# Phosphorylation triggers presynaptic phase separation of Liprin-α3 to control active zone structure

**DOI:** 10.1101/2020.10.28.357574

**Authors:** Javier Emperador-Melero, Man Yan Wong, Shan Shan H. Wang, Giovanni de Nola, Tom Kirchhausen, Pascal S. Kaeser

## Abstract

Liquid-liquid phase separation enables the assembly of membrane-less subcellular compartments, but testing its biological functions has been difficult. The presynaptic active zone, protein machinery in nerve terminals that defines sites for neurotransmitter release, may be organized through phase separation. Here, we discover that the active zone protein Liprin-α3 rapidly and reversibly undergoes phase separation upon phosphorylation by PKC at a single site. RIM and Munc13 are co-recruited to membrane-attached condensates, and phospho-specific antibodies establish Liprin-α3 phosphorylation in vivo. At synapses of newly generated Liprin-α2/α3 double knockout mice, RIM, Munc13 and the pool of releasable vesicles were reduced. Re-expression of Liprin-α3 restored these defects, but mutating the Liprin-α3 phosphorylation site to abolish phase condensation prevented rescue. Finally, PKC activation acutely increased RIM, Munc13 and neurotransmitter release, which depended on the presence of phosphorylatable Liprin-α3. We conclude that Liprin-α3 phosphorylation rapidly triggers presynaptic phase separation to modulate active zone structure and function.

## Introduction

Membrane-free subcellular compartments form through liquid-liquid phase separation, a process in which multivalent, low affinity interactions enable de-mixing of proteins into liquid condensates ^1–3^. These condensates maintain high local protein concentrations and create a plastic environment that enables molecular re-arrangement and exchange with the environment. Compelling work has established that protein complexes for many processes, ranging from gene transcription to neurodegeneration, can be organized as phase condensates, but it has remained challenging to establish which condensates form in vivo and to determine how phase separation controls intracellular functions.

This is particularly true for synaptic transmission. Within a synapse, neurotransmitter release is restricted to specialized presynaptic structures called active zones ^4, 5^. These membrane-attached, dense scaffolds are formed by the multidomain proteins RIM, Munc13, RIM-BP, Piccolo/Bassoon, ELKS and Liprin-α, and are essential for the sub-millisecond precision of synaptic vesicle exocytosis. While several mechanisms of these proteins in release are established, it remains largely unknown how these dense scaffolds assemble, and how they remain dynamic to maintain the high spatiotemporal demands of presynaptic vesicle traffic. Purified RIM1 and RIM-BP2 form liquid condensates in vitro, indicating that active zones may assemble following phase transition principles ^6^, and other subsynaptic compartments may also be organized by phase separation ^7, 8^. Whether phase separation occurs at synapses in vivo, however, remains debated, and whether it is important for controlling hallmark properties of synaptic release, for example its speed and plasticity, is unclear.

Liprin-α proteins have received particular attention as assembly molecules because they control presynaptic structure of invertebrate synapses ^9–12^. They contain N-terminal coiled-coils with Liprin-α homology (LH) regions and three C-terminal SAM domains ^5, 10, 13, 14^. Mammals have four genes (*Ppfia1*-*Ppfia4*) that encode Liprin-α1 to Liprin-α4 ^13^, of which only Liprin-α2 and Liprin-α3 are strongly expressed in the brain and co-localize with active zone markers ^15, 16^. shRNA knockdown of Liprin-α2 ^17^ or genetic deletion of the Liprin-α3 ^16^ causes loss of presynaptic proteins, similar to assembly defects after ablation of the single invertebrate gene ^9–12, 18^. While these data implicate Liprin-α in active zone assembly, the vertebrate Liprin-α functions and their underlying mechanisms are not clear. Liprin-α2 and Liprin-α3 localize normally after genetic disruption of vertebrate active zones ^16, 19, 20^, which may reflect an upstream assembly function similar to invertebrates ^11, 12^, or suggest that Liprin-α proteins are not part of the same protein complex. An upstream function aligns well with the broad interaction repertoire of Liprin-α, which includes active zone proteins, motors, cell adhesion proteins and cytoskeletal elements ^5, 12, 13, 21–25^. Liprin-α interactions are further regulated by phosphorylation ^26^, making it a candidate effector of kinase pathways that control exocytosis, for example of protein kinase A (PKA), phospholipase C (PLC)/protein kinase C (PKC), or Ca^2+^/calmodulin-dependent kinase II (CaMKII) signaling ^27^. In aggregate, previous data suggest that Liprin-α may connect active zone assembly to upstream pathways for synapse development and plasticity.

We here find that PKC phosphorylation of serine-760 (S760) of Liprin-α3 rapidly triggers Liprin-α3 phase separation. RIM and Munc13-1, two important active zone proteins, are co-recruited into plasma-membrane attached phase condensates, reminiscent of active zone assembly. Newly developed double knockout of Liprin-α2 and Liprin-α3 leads to loss of RIM and Munc13-1, impaired vesicle docking and a decreased pool of readily releasable vesicles. Abolishing Liprin-α3 phosphorylation via a single point mutation prevents its phase separation and its ability to reverse defects in active zone structure and in the pool of releasable vesicles. Similarly, we discover rapid enhancement of RIM and Munc13-1 levels at the active zone upon activation of PKC, which necessitates Liprin-α3 phosphorylation. We conclude that active zone structure is dynamically modulated by Liprin-α3 phase condensation under the control of PKC, establishing a role for liquid-liquid phase separation in presynaptic architecture and plasticity.

## Results

### Liprin-α3 rapidly undergoes phase separation under the control of PLC/PKC signaling

Because Liprin-α3 is regulated by phosphorylation and controls active zone assembly ^10, 16, 26^, we asked whether Liprin-α3 is modulated by kinase pathways to control release site structure. Prominent presynaptic pathways operate via PKA, PLC/PKC, and CaMKII signaling ^27^. We expressed mVenus-tagged Liprin-α3 in HEK293T cells and investigated whether activation or inhibition of these pathways alters Liprin-α3 distribution. Under basal conditions, mVenus-Liprin-α3 is predominantly soluble. Strikingly, after addition of the diacylglycerol analogue phorbol 12-myristate 13-acetate (PMA), Liprin-α3 rapidly formed spherical condensates (Figs. 1a, Extended Data Fig. 1a and Movie 1). PMA mimics PLC-induced generation of diacylglycerol and activates PKC, suggesting that Liprin-α3 may be phosphorylated by PKC. This effect was not observed for other manipulations, including inhibiting PKC, or activation or inhibition of PKA or CaMKII. The reorganization of Liprin-α3 into droplets occurred in all cells within minutes, was reversible upon washout, and droplet formation was independent of the mVenus-tag (Figs. 1b, 1c, Extended Data Figs. 1a-1c).

**Figure 1.**
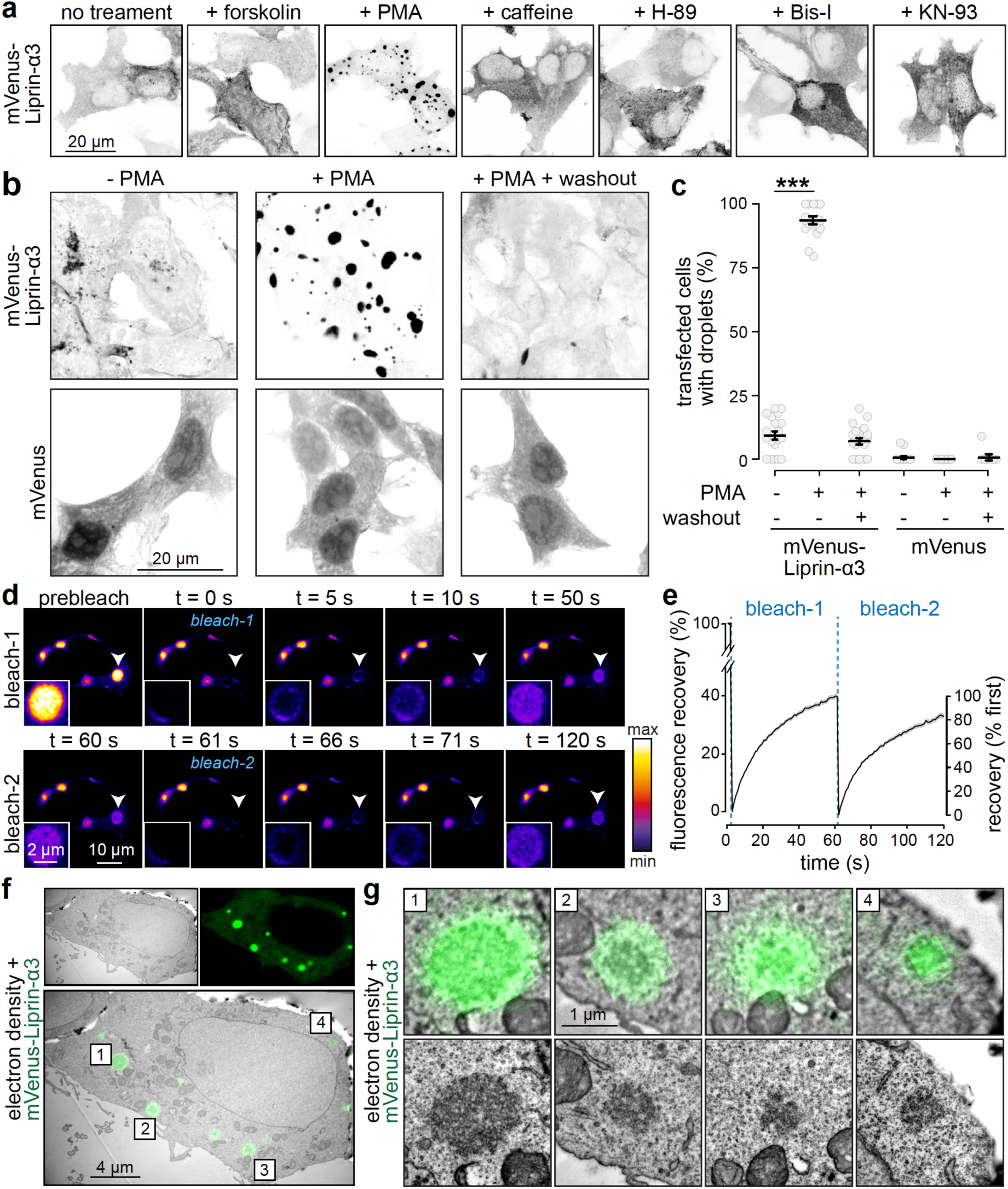
Liprin-α3 forms electron-dense condensates via liquid-liquid phase separation upon mimicking PLC/PKC activation. **(a)** Confocal images of HEK293T cells transfected with mVenus-Liprin-α3, without treatment or in the presence of forskolin (to activate PKA), PMA (to activate PLC/PKC), caffeine (to activate CamKII), H-89 (to inhibit PKA), bisindolylmaleimide-I (to inhibit PKC), or KN-93 (to inhibit CamKII). **(b, c)** Example confocal images (b) and quantification of the percent cells containing droplets (c) of HEK293T cells transfected with mVenus-Liprin-α3 or mVenus alone. Cells were fixed 15 min after PMA addition or six h after washout. N = 21 images/3 independent batches of cells each. **(d, e)** Example time-lapse images (d) and quantification (e) of the fluorescence recovery after photobleaching (FRAP) of mVenus-Liprin-α3 condensates. Two consecutive bleach steps were applied. N = 30 droplets/3 independent transfections. **(f, g)** Correlative light-electron microscopy (CLEM) example images of a HEK293T cell transfected with mVenus-Liprin-α3 and incubated with PMA showing an overview with multiple condensates (f) and detailed individual droplets (g) magnified from the overview image (top) and independently acquired higher magnification images of the same droplets (bottom). Summary data in c, e and g are mean ± SEM. *** p < 0.001 assessed by Kruskal-Wallis tests with Holm post-hoc comparison against the respective - PMA condition. For a time course of phase separation, phase separation of non-tagged Liprin-α3 and liquid droplet fusion, see Extended Data Fig. 1 and Movie 1.

Formation of spherical droplets is indicative of liquid-liquid phase separation ^1, 2^. Principles of liquid dynamics predict droplet fusion, which we observed (Extended Data Figs. 1d, 1e and Movie 1), and exchange of molecules between condensates and the surrounding cytosol. To test exchange, we assessed fluorescence recovery after photobleaching (FRAP), as implemented before to study synaptic liquid phases ^6–8, 28^. Individual condensates recovered to ∼40% of the initial fluorescence at a fast rate (t_1/2 recovery_ < 20 s), and a second bleaching of the same condensates resulted in near-complete recovery, again with t_1/2 recovery_ < 20 s, indicating that the mobile fraction remains fully mobile (Figs. 1d, 1e).

We next assessed whether these fluorescent droplets are indeed membrane-free protein dense condensates. We used correlative light-electron microscopy (CLEM) and found that Liprin-α3 condensates were electron-dense structures without surrounding lipid bilayers (Figs. 1f, 1g). We conclude that Liprin-α3 rapidly and reversibly forms phase-separated condensates as a function of PLC/PKC signaling.

### PKC phosphorylates Liprin-α3 in vitro and in vivo to trigger condensate formation

We hypothesized that PMA triggers PKC activation followed by phosphorylation of Liprin-α3 to induce phase separation. To investigate whether Liprin-α3 is a PKC substrate, we purified GST-fusion proteins covering the entire Liprin-α3 protein, and incubated them with ^32^P-labelled ATP and recombinant PKC (Figs. 2a, 2b). The linker region between the LH and SAM regions most efficiently incorporated ^32^P, and mass spectrometry identified five phosphorylated serine residues (S650, S751, S760, S763 and S764, Extended Data Fig. 2a). Notably, S760, but not other residues, was surrounded by a PKC consensus sequence. To determine whether any of these residues is responsible for phase transition, we engineered point mutations in mVenus-Liprin-α3 to abolish phosphorylation and expressed these constructs in HEK293T cells. S760A and S764A Liprin-α3 were incapable of PMA-induced droplet formation, while other point mutations did not impair it (Extended Data Fig. 2b). To assess whether these residues are phosphorylated, we generated anti-phospho-S760 and -S764 Liprin-α3 antibodies. Both antibodies detected a band at ∼150 KDa in immunoblots of transfected HEK293T cells (Fig. 2c, Extended Data Fig. 2c). Upon PMA addition, and consistent with the PKC consensus sequence, the phospho-S760-Liprin-α3 increased, and disappeared when co-incubated with PKC blockers, while phospho-S764 signals were unchanged. Phospho-S760 Liprin-α3 was not detected in Liprin-α3 knockout neuronal cultures (Fig. 2d), confirming antibody specificity. In vivo, phospho-S760 Liprin-α3 was present in the frontal cortex, hippocampus, cerebellum and brain stem with high perinatal levels that gradually decreased over time (Extended Data Fig. 2d).

**Figure 2.**
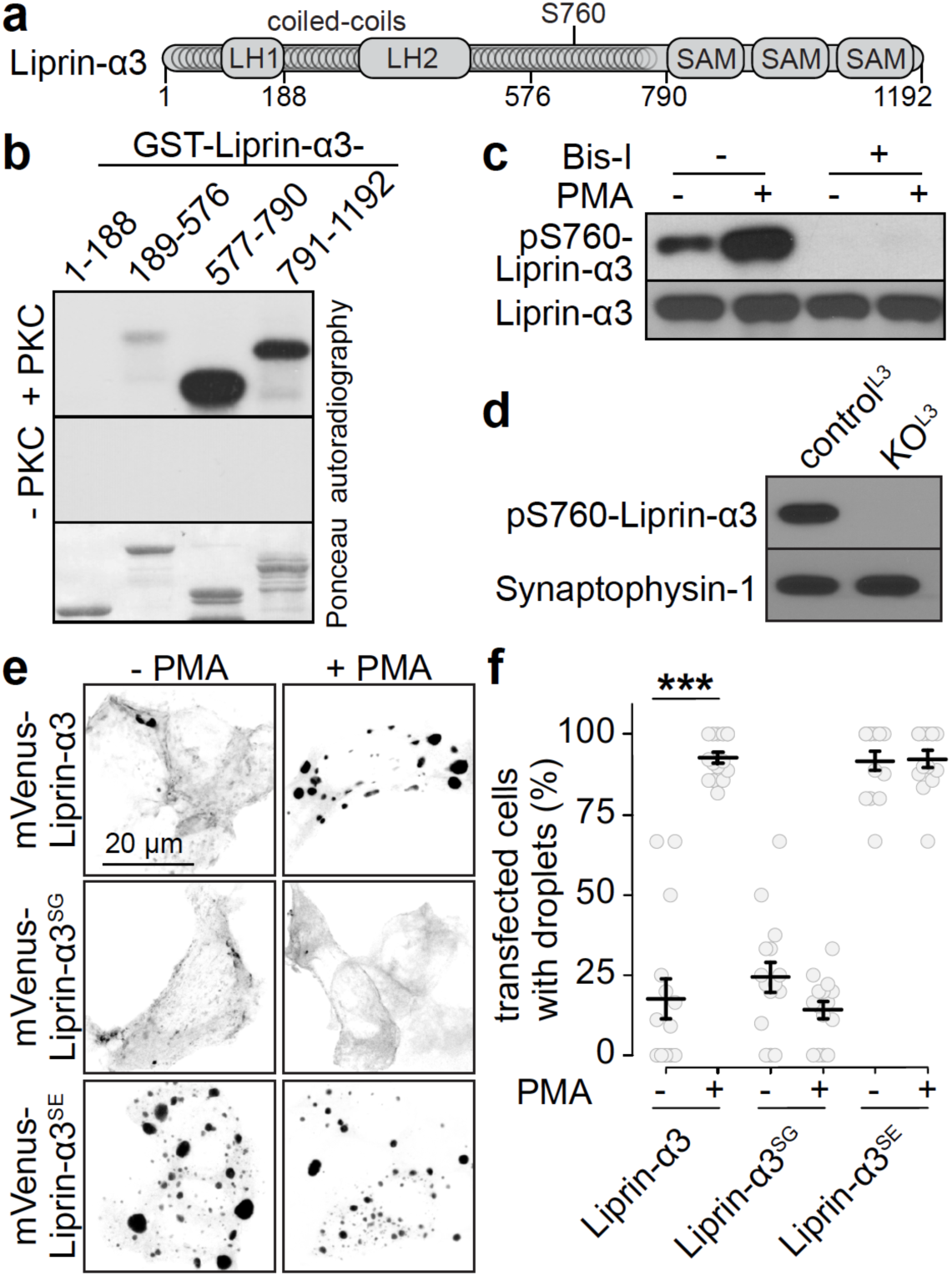
Protein kinase C phosphorylation of Liprin-α3 at serine-760 induces phase separation. **(a)** Schematic of the Liprin-α3 domain structure showing Liprin homology regions 1 and 2 (LH-1 and -2), coiled-coil regions and sterile alpha motifs (SAM). **(b)** Autoradiography (top, middle) and Ponceau staining (bottom) of purified GST-Liprin-α3 fragments incubated with ^32^P-γ-ATP and recombinant PKC (top), or without PKC (middle). **(c)** Western blot of lysates of transfected HEK293T cells expressing Liprin-α3 and incubated with PMA and, where indicated, the PKC inhibitor bisindolylmaleimide-I (Bis-I), and immunoblotted with newly generated anti-phospho-S760 Liprin-α3 or Liprin-α3 antibodies. **(d)** Western blot of lysates of cultured hippocampal neurons from Liprin-α3 knockout mice (KO^L3^) or from heterozygote control mice (control^L3^), with phospho-S760 Liprin-α3 antibodies. **(e, f)** Example confocal images **(e)** and quantification **(f)** of droplet formation in HEK293T cells expressing mVenus-tagged wild type Liprin-α3, phospho-dead Liprin-α3 S760G (Liprin-α3^SG^) or phospho-mimetic Liprin-α3 S760E (Liprin-α3^SE^). N = 15 images/3 independent transfections each. Data in f are mean ± SEM. *** p < 0.001 assessed by Kruskal-Wallis tests. For evaluation of additional potential phosphorylation sites, expression profile of phospho-S760 Liprin-α3 across brain areas and development, and droplet formation of other Liprin-α isoforms, see Extended Data Fig. 2.

Our data establish that PKC phosphorylates Liprin-α3 at S760. To corroborate that this site mediates phase separation, we generated phospho-dead (S760G, Liprin-α3^SG^, using S->G substitution to make it similar to other Liprin-α proteins, Extended Data Fig. 2e) and phospho-mimetic (S760E, Liprin-α3^SE^) mutants. Liprin-α3^SG^ abolished PKC-induced phase separation, and Liprin-α3^SE^ formed constitutive condensates independent of PKC activation (Figs. 2e, 2f). S760 is conserved in Liprin-α3 across vertebrates, but it is a glycine residue in the other three vertebrate Liprin-α’s and in invertebrate proteins (Extended Data Fig. 2e). In line with a Liprin-α3-specific function of S760, mimicking PLC/PKC signaling in HEK293T cells expressing Liprin-α1, -α2 or -α4 did not change the distribution of any of these proteins, which was predominantly soluble for Liprin-α1 and -α4, and droplet-like for Liprin-α2 (Extended Data Figs. 2f, 2g). We conclude that PKC phosphorylates S760 of Liprin-α3 in vitro and in vivo to trigger Liprin-α3 phase separation.

### Liprin-α3, RIM1α and Munc13-1 are co-recruited into membrane-attached liquid condensates

We reasoned that if phase separation of Liprin-α3 controls active zone assembly, active zone proteins must interact with this liquid phase. Co-expression of cerulean-Liprin-α3 with either RIM1α-mVenus or Munc13-1-tdTomato in HEK293T cells resulted in recruitment of each protein into PMA-induced condensates (Extended Data Fig. 3a). Discrete, PMA-insensitive condensates were also observed when RIM1α was expressed alone (Extended Data Fig. 3b), in agreement with its intrinsic ability to phase separate ^6^. Munc13-1 did not form droplets on its own, but PMA-dependent membrane recruitment was observed as previously described ^29–31^.

Co-expression of cerulean-Liprin-α3 with both RIM1α-mVenus and Munc13-1-tdTomato in HEK293T cells resulted in large protein condensates, and addition of PMA increased their number and size (Figs. 3a-3c). Remarkably, these condensates were not distributed throughout the cytosol, different from Liprin-α3 phase condensates. Instead, they were in the cell periphery in close proximity to the plasma membrane, and the condensates contained all three proteins. To assess whether they were membrane attached, we used CLEM on PMA-treated cells. The fluorescent signals were highly overlapping with large protein densities that were not enclosed by membranes, but instead appeared attached at one side to the plasma membrane (Figs. 3d, 3e).

**Figure 3.**
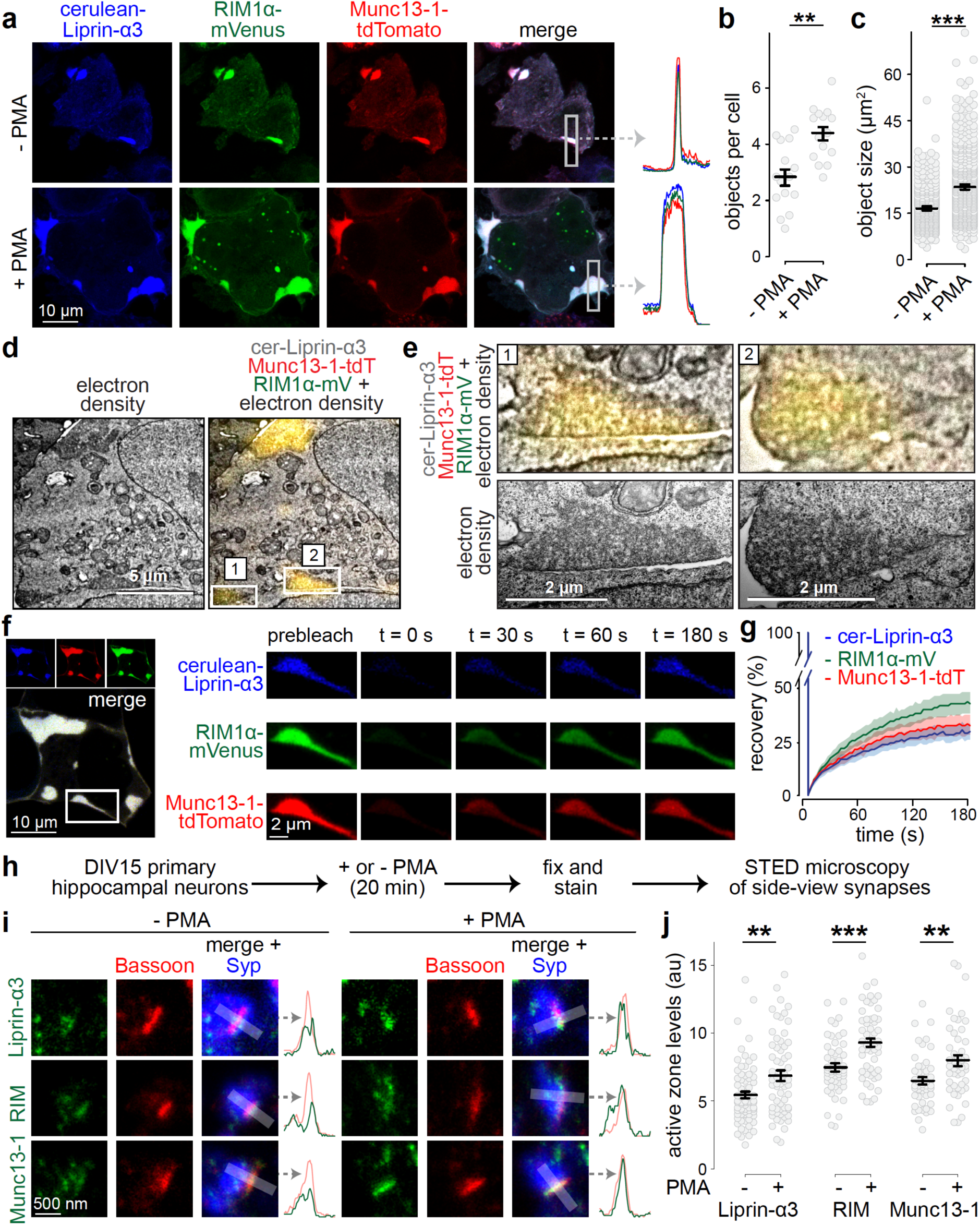
Liprin-α3, Munc13 and RIM are co-recruited into phase condensates at the plasma membrane. **(a-c)** Example confocal images including line profiles of highlighted regions (a) and quantification (b) of phase condensates in HEK293T cells transfected with cerulean-Liprin-α3, RIM1α-mVenus and Munc13-1-tdTomato in the absence or presence of PMA. Quantification of the number (b) and size (c) of protein condensates is shown. N = 15 images/3 independent transfections. **(d, e)** CLEM example images of a HEK293T cell transfected with cerulean-Liprin-α3, RIM1α-mVenus and Munc13-1-tdTomato and incubated with PMA showing an overview with multiple condensates (d) and detailed individual condensates (e) magnified from the overview image (top) and independently acquired images at higher magnification of the same droplets (bottom). **(f, g)** Example of FRAP experiment (f) and quantification (g) of droplets in HEK293T cells transfected with cerulean-Liprin-α3, RIM1α-mVenus and Munc13-1-tdTomato. N = 20 droplets/3 independent transfections. **(h)** Schematic of the assessment of effects of PKC activation on active zone assembly. **(i, j)** Example STED images (i) and quantification (j) of the intensity of endogenous Liprin-α3, RIM and Munc13-1 at the active zone. Synapses in side-view were identified by the active zone marker Bassoon (imaged by STED microscopy) aligned at the edge of a synaptic vesicle cluster marked by Synaptophysin (Syp; imaged by confocal microscopy). An example intensity profile of the intensity of the protein of interest and Bassoon is shown on the right of each image set. Peak intensities were measured in intensity profiles and plotted in j. Liprin-α3: N = 71 synapses/3 independent cultures (- PMA) and 63/3 (+ PMA); RIM: N = 55/3 (- PMA) and 54/3 (+ PMA); Munc13-1 N = 46/3 (- PMA) and 44/3 (+ PMA). Data are shown as mean ± SEM. ** p < 0.01, *** p < 0.001 assessed by Mann-Whitney rank sum tests in b, c and j. For assessment of single and double transfections, and FRAP without PMA treatment see Extended Data Fig. 3, for a description of STED analyses and peak positions for each protein, see Extended Data Fig. 4.

We finally used FRAP to assess turnover of Liprin-α3, RIM1α and Munc13-1 in these condensates. All three proteins rapidly recovered when the entire condensate was bleached (Figs. 3f, 3g) or when only small areas within large condensates were bleached (Extended Data Fig. 3c). Hence, membrane-attached condensates containing Liprin-α3, RIM1α and Munc13-1 follow liquid dynamics. Overall, these data establish that Liprin-α3, RIM1α and Munc13-1 co-exist in protein-dense liquid condensates attached to the plasma membrane, and formation of these condensates is enhanced by PLC/PKC signaling.

### PLC/PKC signaling increases active zone levels of Liprin-α3, RIM and Munc13-1 at synapses

Our findings suggest that activating PKC induces the formation of active zone-like, membrane-bound liquid condensates in transfected cells. If physiologically relevant, activation of this pathway should result in changes in active zone protein complexes at synapses. To test this, we assessed active zone levels of endogenous Liprin-α3, RIM and Munc13-1 at synapses of cultured hippocampal neurons using stimulated emission depletion (STED) microscopy (Fig. 3h-3j). As described previously ^16, 32–34^, we restricted the analysis to side-view synapses to avoid skewing results by synapse orientation. Side-view synapses were identified by the position of a bar-shaped active zone (marked by Bassoon, imaged in STED mode) relative to a synaptic vesicle cloud (identified by Synaptophysin, imaged in confocal mode), and the peak levels of proteins at active zones were measured within 100 nm of the Bassoon peak (see Extended Data Fig. 4a for an outline of synapse selection and analyses). Liprin-α3, RIM and Munc13-1 were predominantly clustered at the active zone with peak intensities falling within 50 nm from the peak of Bassoon (Extended Data Fig. 4b) as shown before ^16^. Addition of PMA produced a significant 20-30% increase in peak active zone levels of Liprin-α3, RIM and Munc13-1 (Figs. 3i, 3j) without affecting Bassoon (Extended Data Figs. 4c-4e). Hence, mimicking PLC/PKC activation induces structural active zone plasticity with enhanced recruitment of RIM, Munc13-1 and Liprin-α3.

### Knockout of Liprin-α2 and Liprin-α3 alters presynaptic composition and ultrastructure

If Liprin-α phase separation controls active zone assembly, Liprin-α knockout should impair its structure and function. We generated new knockout mice to simultaneously ablate Liprin-α2 and Liprin-α3, the main synaptic Liprin-α proteins (Extended Data Fig. 5) ^16, 35^. Conditional Liprin-α2 knockout mice (Liprin-α2^f/f^), generated by homologous recombination with exon 14 flanked by loxP sites (Extended Data Figs. 6a-6e), were crossed to homozygosity and subsequently bred to previously generated constitutive Liprin-α3 knockout mice (Liprin-α3^-/-^) ^16^ (Fig. 4a). We used cultured hippocampal neurons of Liprin-α2^f/f^/Liprin-α3^-/-^ mice infected with lentivirus expressing cre recombinase (to generate KO^L23^ neurons) and neurons from Liprin-α2^f/f^/Liprin-α3^+/-^ mice infected with lentiviruses that express truncated, inactive cre recombinase (to generate control^L23^ neurons). First, we assessed the composition of synapses by confocal microscopy by measuring protein levels within synapses (Fig. 4b). Liprin-α2 and Liprin-α3 were efficiently removed and the remaining signals are typical for antibody background ^19, 33^. The levels of RIM and Munc13-1 were decreased by 25–40%, without significant changes in Bassoon, RIM-BP2, and other synaptic proteins. Surprisingly, the synaptic levels of Ca_V_2.1 were increased by ∼50%, as were those of Synapsin-1, but Synaptophysin levels were unchanged (Figs. 4b,4c, Extended Data Figs. 6f-6h).

**Figure 4.**
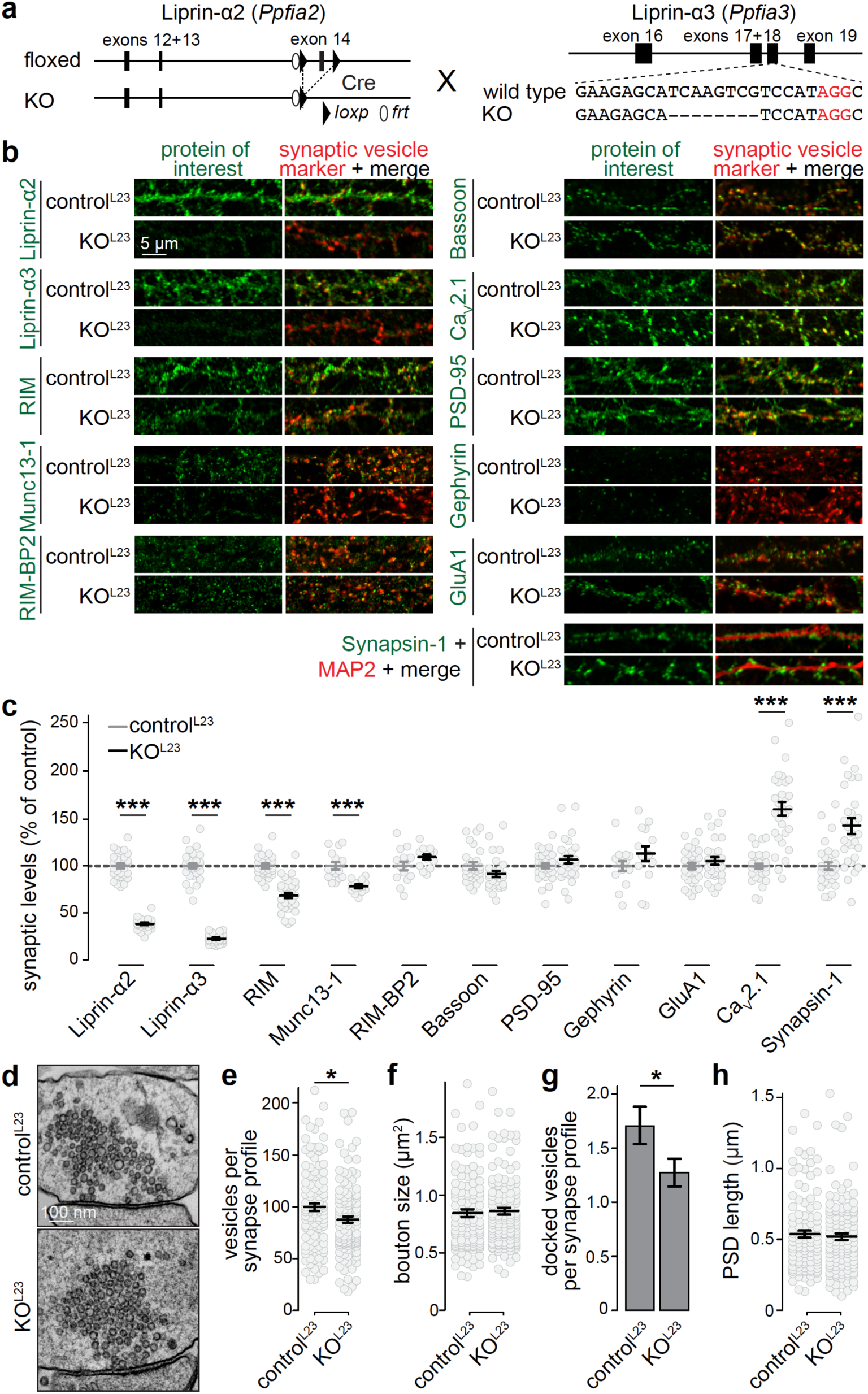
Double Liprin-α2/α3 knockout alters presynaptic composition and ultrastructure. **(a)** Schematic for simultaneous knockout of Liprin-α2 and -α3. Newly generated mice in which Liprin-α2 can be removed by cre recombination (Liprin-α2^f/f^) were crossed to previously published constitutive Liprin-α3 knockout (Liprin-α3^-/-^) mice that were generated by CRISPR/Cas9-mediated genome editing (deleted sequence is represented with dashes) ^16^. Cultured hippocampal neurons of Liprin-α2^f/f^/Liprin-α3^-/-^ mice infected with lentivirus expressing cre recombinase were used to generate KO^L23^ neurons, and neurons from Liprin-α2^f/f^ x Liprin-α3^+/-^ mice infected with lentiviruses that express truncated, inactive cre recombinase were used to generate control^L23^ neurons. **(b, c)** Example confocal images (b) and quantification (c) of neurons immunostained for either Liprin-α2, Liprin-α3, RIM, Munc13-1, RIM-BP2, Bassoon, Ca_V_2.1, PSD-95, Gephyrin or GluA1 and Synapsin (for Munc13-1, RIM-BP2 and Gephyrin) or Synaptophysin (all others) as vesicle marker, or Synapsin-1 and MAP2. Quantification in c was performed in regions of interest (ROIs) defined by the synaptic vesicle marker and normalized to the average control^L23^ levels per culture. N = 30 images/3 independent cultures per genotype per protein of interest, except for Munc13-1 and RIM-BP2 (N = 15/3) and Gephyrin (N = 14/3). **(d-h)** Example electron micrographs of synapses (d) and quantification of the number of synaptic vesicles per section (e), bouton size (f), number of docked vesicles per section (g) and PSD length (h) of neurons fixed by high-pressure freezing followed by freeze substitution, control^L23^: N = 111 synapses/2 independent cultures, KO^L23^: N = 123/2. All data are mean ± SEM and were analyzed using Mann-Whitney U tests, except for PSD-95 and Gephyrin in c, for which t-tests were used. * p < 0.05, *** p < 0.001. For synaptic localization of Liprin-α1-α4 see Extended Data Fig. 5, generation of Liprin-α2^f/f^ mice and analysis of Synaptophysin levels of the experiments shown in c see Extended Data Fig. 6, and for STED analysis of active zone localization of RIM, Munc13-1, RIM-BP2 and Ca_V_2.1 in KO^L23^ neurons, see Extended Data Fig. 7.

We next asked whether these changes in protein levels occur at the active zone and whether they are present in both excitatory and inhibitory synapses (marked with PSD-95 and Gephyrin, respectively (Extended Data Fig. 7)). At side-view synapses, the peak levels of Munc13-1, RIM, RIM-BP2 and Ca_V_2.1 peaked at ∼ 100 nm from the postsynaptic markers in agreement with their active zone localization ^16, 34^. Decreased active zone levels of Munc13-1 and RIM were observed in both synapse types in KO^L23^ neurons, while the increase of Ca_V_2.1 was restricted to excitatory synapses. Hence, Liprin-α2 and -α3 are necessary to maintain normal active zone structure.

High-pressure freezing followed by freeze substitution and electron microscopic imaging was used to investigate synaptic ultrastructure. The number of synaptic vesicles per synapse profile was decreased by ∼15% in KO^L23^ synapses, without changes in the overall bouton size or postsynaptic densities (Figs. 4d-h). A ∼25% reduction of docked vesicles (identified as vesicles with no detectable space between the electron-dense vesicular and target membranes) was observed upon Liprin-α2/α3 knockout, consistent with a partial loss of the docking proteins RIM and Munc13-1. We conclude that Liprin-α2 and Liprin-α3 are involved in maintaining presynaptic ultrastructure, specifically the number of vesicles per bouton and the number of docked vesicles.

### Synapse-specific impairments in neurotransmitter release at KO^L23^ synapses

The altered levels of Munc13-1, RIM and Ca_V_2.1 and the decreased docking predict changes in synaptic secretion. Indeed, in whole-cell electrophysiological recordings, the frequency of spontaneous miniature excitatory and inhibitory postsynaptic currents (mEPSCs and mIPSCs, respectively) was decreased in KO^L23^ neurons, but their amplitudes were unchanged (Figs. 5a-5c, 5j-5l), establishing presynaptic roles of Liprin-α in synaptic vesicle release.

**Figure 5.**
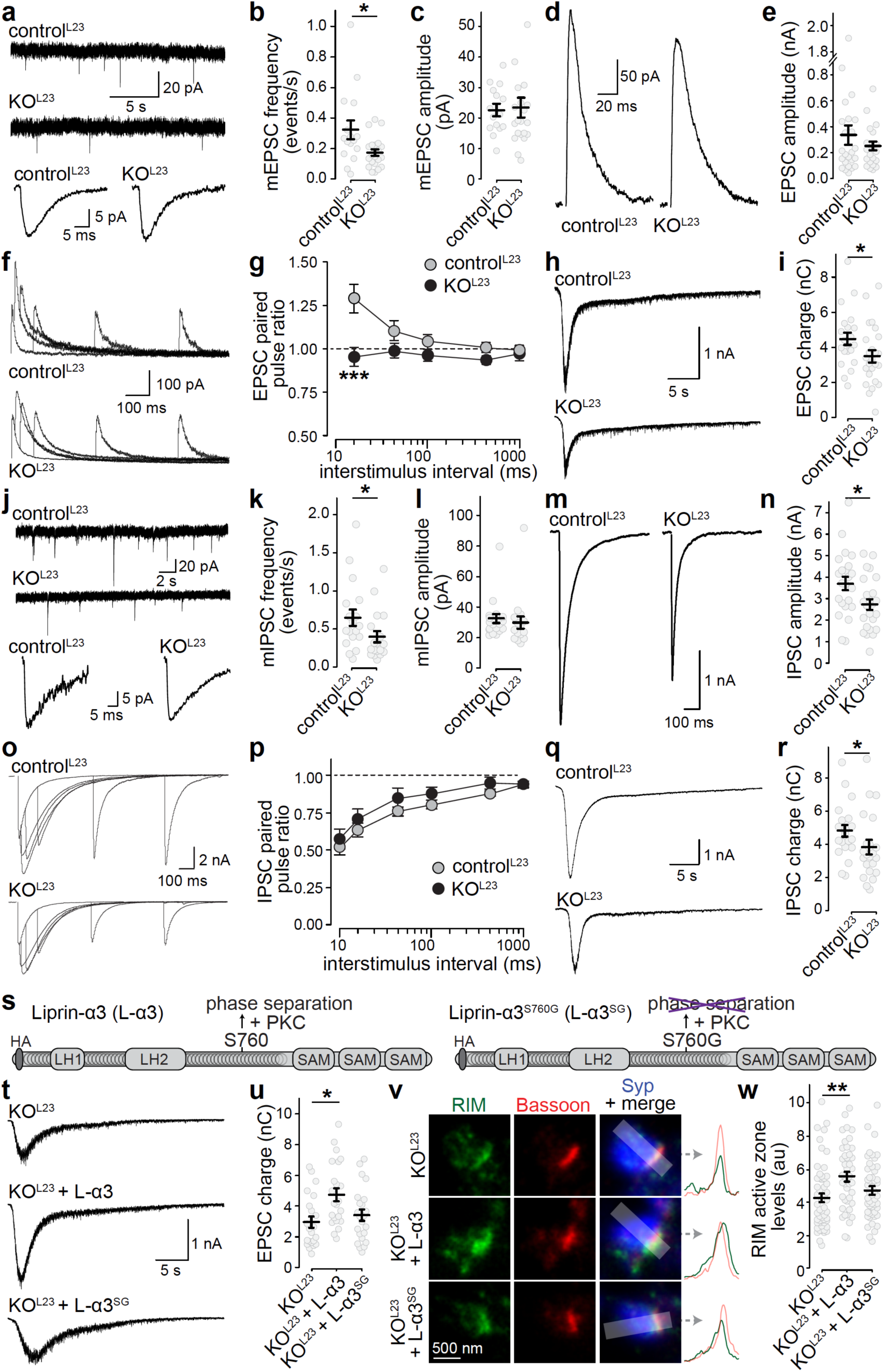
Liprin-α2/α3 double knockout impairs neurotransmitter release. **(a-c)** Example traces (a) of spontaneous miniature excitatory postsynaptic currents (mEPSC) recordings (top) and an averaged mEPSC of a single cell (bottom) and quantification of mEPSC frequency (b) and amplitude (c). For mEPSC frequency, control^L23^: N = 16 cells/3 independent cultures, KO^L23^: N = 21/3. For amplitude, control^L23^: N = 15/3, KO^L23^: N = 20/3. **(d, e)** Example traces (d) and average amplitudes (e) of single action potential-evoked NMDA receptor-mediated EPSCs, control^L23^: N = 21/3, KO^L23^: N = 20/3. **(f, g)** Example traces (f) and average NMDA-EPSC paired pulse ratios (g, PPRs) at various interstimulus intervals, control^L23^: N = 21/3, KO^L23^: N = 20/3. **(h, i)** Example traces (h) and quantification (i) of the AMPA receptor-mediated EPSC charge in response to a local 10 s puff of 500 mOsm sucrose to estimate the RRP. control^L23^: N = 21/3, KO^L23^: N = 25/3. **(j-r)** Same as a-i, but for IPSCs, k+l: for mIPSC frequency, control^L23^: N = 18/3, KO^L23^: N = 18/3, for amplitude, control^L23^: N = 21/3, KO^L23^: N = 15/3; m + n: control^L23^: N = 23/3, KO^L23^: N = 24/3; o + p: control^L23^: N = 23/3, KO^L23^: N = 24/3, q + r: control^L23^: N = 22/3, KO^L23^: N = 22/3. **(s)** Diagram of the rescue experiment with Liprin-α3 expression via lentiviral transduction. **(t, u)** Example traces (t) and quantification (u) of sucrose-triggered EPSCs, KO^L23^: N = 22/3, KO^L23^ + Liprin-α3 (L-α3): N = 23/3, KO^L23^ + Liprin-α3^S760G^ (L-α3^SG^): N = 23/3. **(v, w)** Representative STED images (v) and quantification (w) of RIM at the active zone of side-view synapses as in Figs. 3i-3j. KO^L23^: N = 56 synapses/3 independent cultures, KO^L23^ + L-α3: N = 43/3, KO^L23^ + L-α3^SG^: N = 50/3. All data are mean ± SEM, * p < 0.05, ** p < 0.01 analyzed using Mann-Whitney U tests (b, c, i, k, l), t-tests (e, n, r), two-way ANOVA (g, p), or Kruskal-Wallis (u, w) with Tukey-kramer (g, p) or Holm (u, w) post-hoc comparison against KO^L23^. For a direct comparison of control^L23^, KO^L23^ and KO^L23^ + L-α3 and for STED localization of rescue Liprin-α, see extended Data Fig. 8.

We used electrical stimulation or stimulation by hyperosmotic sucrose to evoke EPSCs (Figs. 5d-5i) and IPSCs (Figs. 5m-5r). Release evoked by an action potential is proportional to the product of the number of vesicles that can be released (readily releasable pool, RRP) and the likelihood of a vesicle to be released (vesicular release probability, p) ^36, 37^. For action-potential triggered EPSCs, NMDA-receptor currents were measured instead of AMPA-receptor currents to avoid network activity that is prominent when AMPA-receptors are not blocked. Similar to confocal and STED microscopy, we observed synapse-specific changes. At excitatory and inhibitory KO^L23^ synapses, the RRP estimated by the application of hypertonic sucrose was decreased (Figs. 5h, 5i, 5q, 5r), quantitatively matching the reduction in Munc13, RIM and docked vesicles (Fig. 4, Extended Data Fig. 7). We estimated p by measuring paired pulse ratios (PPRs), where the response ratio of two consecutive pulses at short interstimulus intervals is inversely correlated with p ^36^. Excitatory KO^L23^ synapses had an increased p (Figs. 5f, 5g), matching well with the increased presence of Ca^2+^ channels (Fig. 4, Extended Data Fig. 7). Together, the reduction in RRP and increase in p offset one another and led to a normal EPSC (Fig. 5d, 5e). In contrast, p was unaffected at inhibitory synapses (Figs. 5o, 5p), matching with normal Ca^2+^ channel levels, and leading to an overall decrease in the IPSC amplitude due to the RRP decrease (Figs. 5m, 5n). In summary, the electrophysiological phenotypes match with the structural active zone effects. Knockout of Liprin-α2/α3 leads to reduction in docking, protein machinery for docking and priming and the pool of releasable vesicles at excitatory and inhibitory synapses, and a select increase in Ca^2+^ channels and release probability at excitatory synapses.

### PKC phosphorylation of Liprin-α3 at S760 enhances the readily releasable pool

Next, we asked whether re-expression of wild type Liprin-α3 reverses the presynaptic phenotypes of KO^L23^ neurons. Lentiviral expression of Liprin-α3 in KO^L23^ neurons restored active zone levels of Liprin-α3 (Extended Data Figs. 8a-8d), the RRP (Extended Data Figs. 8e, 8f), as well as the reduced active zone levels of RIM (Extended Data Figs. 8g, 8h). Hence, the active zone impairments at excitatory KO^L23^ synapses are reversible by re-expression of Liprin-α3.

We reasoned that if Liprin-α3 functions depend on its propensity to phase separate, its ability to rescue should be altered when S760 is mutated to abolish phase separation. We directly compared the ability of wild type Liprin-α3 and Liprin-α3^SG^ to rescue RRP and RIM (Fig. 5s-5w). Liprin-α3 proteins (N-terminally tagged with an HA epitope) were expressed in KO^L23^ neurons using lentiviral transduction. At DIV15, both forms of Liprin-α3 were enriched at active zones, but the peak levels of Liprin-α3^SG^ were somewhat decreased compared to Liprin-α3 (∼15%; Extended Data Figs. 8i-8l). Expression of wild type Liprin-α3 increased the RRP and RIM active zone levels by ∼40% in KO^L23^ neurons, while expression of Liprin-α3^SG^ failed to produce any significant increase (Figs. 5t-5w). We conclude that the PKC phosphorylation site of Liprin-α3, which drives phase separation, is essential for a normal RRP and normal active zone structure.

### Liprin-α3 phase separation acutely modulates active zone structure and function

PLC/PKC signaling acutely enhances active zone assembly (Fig. 3) and neurotransmitter release ^38–40^. We hypothesized that this enhancement may be mediated by phosphorylation and phase separation of Liprin-α3, and compared the effect of PKC activation by PMA in KO^L23^ neurons expressing either wild type Liprin-α3 or phase separation-incapable Liprin-α3^SG^. PMA rapidly enhanced mEPSC frequencies and amplitudes (Figs. 6a-6f) as observed before ^40^, indicating that these pathways potentiate synaptic transmission through pre- and postsynaptic effectors. The magnitude of the increase of the mEPSC frequency was impaired by 50% in Liprin-α3^SG^ expressing neurons (Fig. 6a-6c), establishing that Liprin-α phosphorylation is important for this enhancement. Similarly, the RRP estimated by hyperosmotic sucrose application was increased, but this was significantly tempered when non-phosphorylatable Liprin-α3 was present (Fig. 6g-6i). It is noteworthy that the RRP enhancement is overestimated because of the robust increase in mEPSC amplitude (Fig. 6f), and as a consequence the impairment in pool enhancement of Liprin-α3^SG^ is likely underestimated. In summary, these data indicate that PKC phosphorylation and phase separation of Liprin-α3 modulate the RRP.

**Figure 6.**
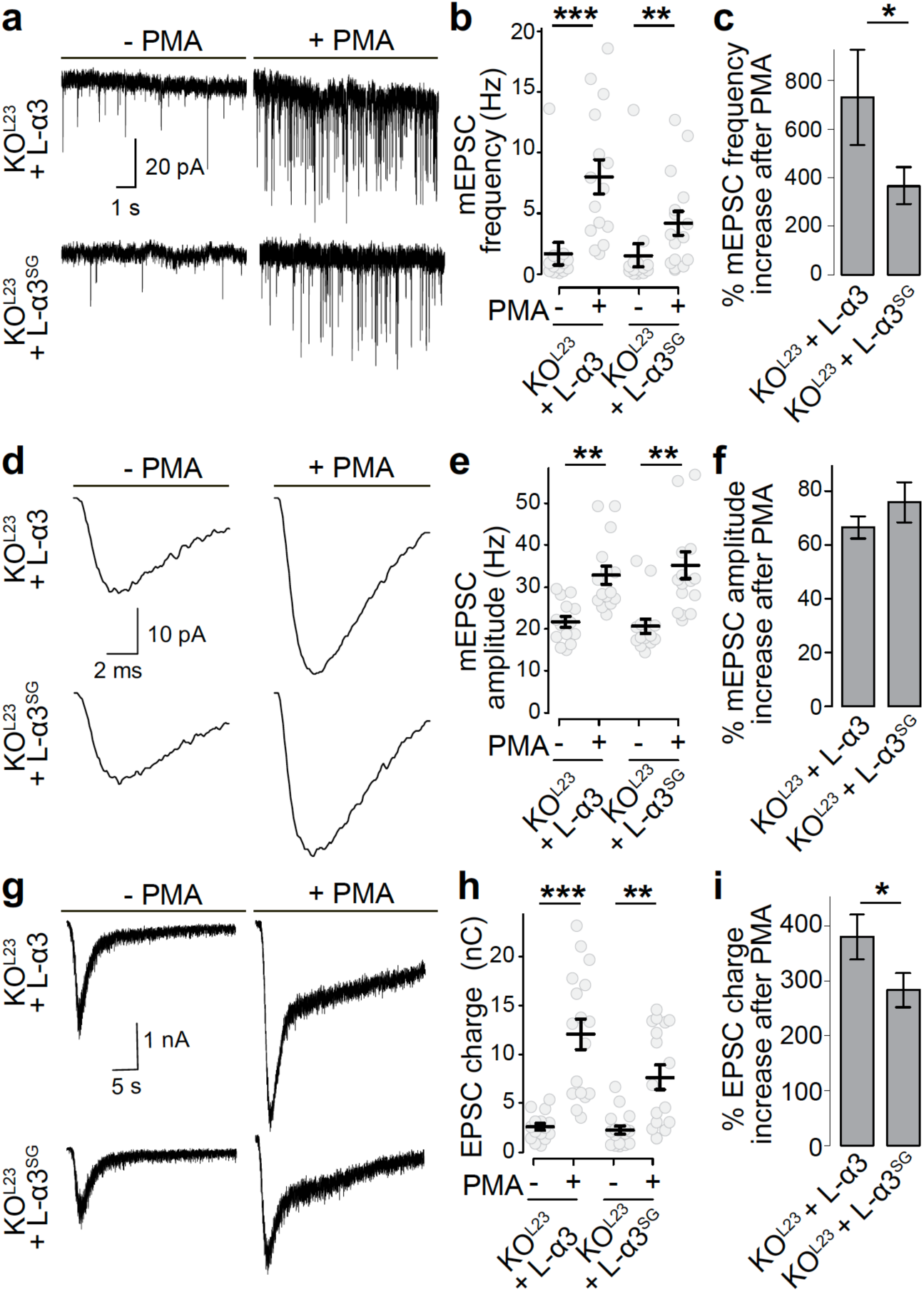
PKC phosphorylation of Liprin-α3 enhances synaptic vesicle release. **(a-c)** Example traces (a) and quantification of mEPSC frequencies (b, c) in KO^L23^ neurons rescued with wild type Liprin-α3 (L-α3) or non-phosphorylatable Liprin-α3 S760G (L-α3^SG^) that does not from phase condensates. The percent increase upon PMA addition over naïve conditions per culture is shown in c. KO^L23^ + L-α3: N = 14 cells/3 independent cultures (- PMA) and 15/3 (+ PMA); KO^L23^ + L-α3^SG^: N = 14/3 (- PMA) and 16/3 (+ PMA). **(d-f)** Average mEPSC from a single cell (d) and quantification of mEPSC amplitudes (e, f). N as in b, c. **(g-i)** Example traces (g) and quantification (h, i) of the EPSC charge in response to a local 10 s puff of 500 mOsm sucrose to estimate the RRP. KO^L23^ + L-α3: N = 15/3 (- PMA) and 17/3 (+ PMA), KO^L23^ + L-α3^SG^: N = 17/3 (- PMA) and 17/3 (+ PMA). All data are mean ± SEM, * p < 0.05, ** p < 0.01, *** p < 0.001 as analyzed by Kruskal-Wallis test and post-hoc (Holm) analysis versus the corresponding - PMA control (b, e, h) or a Mann-Whitney rank sum test (c, f, i).

We finally investigated whether Liprin-α3 phase separation controls active zone structure. We assessed side-view synapses of KO^L23^ neurons, or of KO^L23^ neurons expressing either Liprin-α3 or Liprin-α3^SG^. In both rescue conditions, Liprin-α3, RIM and Munc13-1 were enriched at the active zone. As observed in Fig. 3h-3j, active zone levels of these proteins, but not of Bassoon, robustly increased upon PMA addition by ∼30-35% when Liprin-α3 was present (Fig. 7a-7i, Extended Data Fig. 9). This increase, however, was significantly impaired and indistinguishable from KO^L23^ neurons when only Liprin-α3^SG^ was present. Together, these data show that active zone structure is rapidly modulated by PLC/PKC signaling via phosphorylation and phase separation of Liprin-α3.

**Figure 7.**
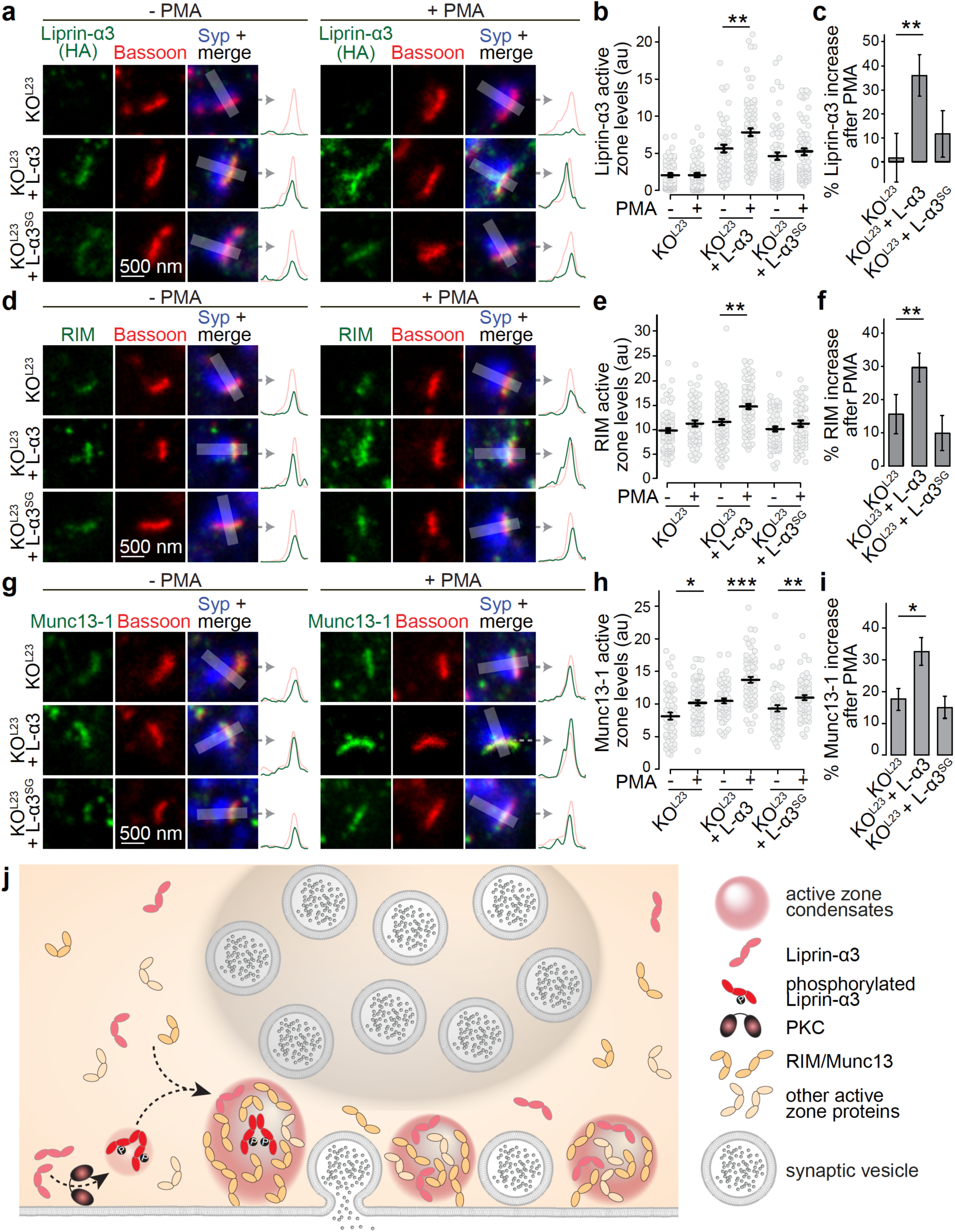
The PKC phosphorylation site of Liprin-α3 acutely modulates active zone assembly. **(a-c)** Example STED images and their intensity profiles (a) and quantification (b, c) of Liprin-α3 (detected by anti-HA antibodies) in side-view synapses in the presence or absence of PMA. The increase upon PMA addition normalized to corresponding - PMA controls is shown in c, KO^L23^: N = 70 synapses/3 cultures (- PMA) and 71/3 (+ PMA), KO^L23^ + L-α3: N = 54/3 (- PMA) and 83/3 (+ PMA); KO^L23^ + L-α3^SG^: N = 63/3 (- PMA) and 73/3 (+ PMA). **(d-i)** Experiments as shown in in a-c, but for RIM (d-f) and Munc13-1 (g-h). RIM (d-f): KO^L23^: N = 81/3 (- PMA) and 61/3 (+ PMA), KO^L23^ + L-α3: N = 75/3 (- PMA) and 84/3 (+ PMA); KO^L23^ + L- α3^SG^: N = 65/3 (- PMA) and 59/3 (+ PMA). Munc13-1 (g-h): KO^L23^: N = 54/3 (- PMA) and 67/3 (+ PMA), KO^L23^ + L-α3: N = 55 /3 (- PMA) and 67/3 (+ PMA), KO^L23^ + L-α3^SG^: N = 57/3 (- PMA) and 62/3 (+ PMA). **(j)** Working model for the control of active zone structure through phase separation of Liprin-α3. The formation of phase condensates is triggered by PKC phosphorylation of Liprin-α3 at serine- 760 and Munc13-1 and RIM are recruited into these release site condensates for boosting neurotransmitter secretion. Data are shown as mean ± SEM, * p < 0.05, ** p < 0.01, *** p < 0.001 as analyzed by Kruskal-Willis tests and post-hoc analysis (Holm) versus the corresponding - PMA control (b, e and h) or versus KO^L23^ (c, f, and i).

## Discussion

Self-assembly of proteins into liquid phases is a biophysical mechanism used by cells for the formation of membrane-less compartments ^1, 2^. We investigated molecular pathways that drive and modulate assembly of the presynaptic active zone, and demonstrate that (1) PKC phosphorylates Liprin-α3 at S760 to drive the formation of membrane-attached liquid condensates containing RIM1α and Munc13-1, (2) genetic ablation of the synaptic Liprin-α proteins leads to defects in active zone structure and function, including the loss of RIM and Munc13-1, and (3) RIM and Munc13-1 active zone levels and neurotransmitter release are acutely upregulated by PKC phosphorylation of S760 followed by phase separation of Liprin-α3. These results lead to a model in which presynaptic phase separation triggered by Liprin-α3 phosphorylation rapidly induces plasticity in active zone structure and neurotransmitter release (Fig. 7j).

### Phase separation of Liprin-α3

Our work establishes a fast mechanism that triggers phase separation of Liprin-α3 into liquid condensates via phosphorylation at S760. Phosphorylation of this region between the LH and SAM domains likely leads to the formation of condensates by increasing Liprin-monomer self-assembly past a critical threshold. This increase could be mediated by enhancing or enabling interactions of the phosphorylated linker itself, by recruitment of adaptors, or by inducing Liprin-α3 conformational changes that expose previously occluded domains to enable new Liprin-α3 interactions. The third scenario appears most likely because S760 is not part of the N-terminal sequences that mediate Liprin-α dimerization ^13, 14^ and in which a gain-of-function mutant that promotes active zone assembly was isolated ^12^.

There are notable differences in condensate formation across vertebrate Liprin-α proteins. Only Liprin-α2 and Liprin-α3 phase separate, in line with their competition for positioning at the active zone ^16^. However, roles in active zone structural plasticity are likely unique to Liprin-α3 because PKC-mediated triggering of phase separation is limited to this isoform. Liprin-α1 and Liprin-α4 do not form condensates in transfected cells and may either operate though different mechanisms or lack important components for phase condensation in these cells. Importantly, these Liprins show a prominent dendritic localization ^15, 41^ and at least Liprin-α1 operates in neuronal arborization ^42^. Together, a picture emerges where the ability to phase separate determines the cellular function of Liprin-α.

### Interactions of presynaptic liquid phases

RIM1α and RIM-BP2 form liquid condensates in vitro, and these condensates organize tethering of voltage-gated calcium channels ^6, 43^. The question arises whether Liprin-α3 is part of the same phase within a nerve terminal, or whether multiple independent phases co-exist. The current evidence is most compatible with a model of multiple distinct phases. First, active zone levels of RIM and Munc13-1 decrease upon ablation of Liprin-α2 and Liprin-α3, but those of Ca_V_2.1 increase and those of RIM-BP2 are unchanged. Similarly, ablation of RIM-BP and RIM ^20^, or of RIM and ELKS ^19^, does not lead to loss of presynaptic Liprin-α. Hence, these proteins are at least partially in distinct protein complexes, or phases. Second, Ca_V_2.1 and Munc13-1 do not co-localize when assessed at nanometer resolution using immunogold labeling, indicating that distinct clustering mechanisms are present ^44^. In aggregate, it appears most likely that distinct liquid assemblies may exist within an active zone, one containing RIM1 and RIM-BP2 to tether calcium channels ^3, 6^, and a different phase with Liprin-α, RIM and Munc13-1. It is interesting that RIM may participate in multiple condensates, perhaps suggesting that it promotes interactions across liquid phases. This may also be true for synaptic vesicle clusters, which are organized through Synapsin phase separation ^7^. Release requires the transition of synaptic vesicles from the cluster to release sites. It appears possible that RIM/Liprin-α/Munc13 phases embody such sites (this study and ^45, 46^), and that RIM allows for recruitment of vesicles from the vesicle phase to release sites, consistent with its roles in vesicle docking ^43, 47, 48^. As such, the tethering and docking reaction could be seen as the transition of a vesicle from Synapsin-phase association to active zone-phase association.

The existence of multiple phase separation-based pathways for active zone assembly may explain difficulties in understanding its assembly mechanisms. Removal of each protein family, for example of Liprin-α (this study), RIM ^43, 49, 50^, or RIM-BP ^51, 52^, leads to at most partial assembly defects, but combinations of mutations are required to disrupt active zones ^19, 20, 53^. This redundancy may also be the reason why active zone protein deletions can lead to synapse-specific secretory deficits ^54, 55^. In summary, this and previous work support the model that there is no single master active zone organizer. Instead, redundant low-affinity interactions organize release sites ^4^. An important remaining question is how these phases are attached to the target membrane. Because disruption of the predominant candidate mechanisms did not lead to active zone disassembly or displacement from the target membrane ^33, 34, 56, 57^, these mechanisms remain obscure.

### Presynaptic phase separation in active zone assembly and function

Previously described active zone condensates form constitutively ^6^, but modulating their formation is ideally suited to explain rapid changes during plasticity. Phosphorylation has been found to regulate phase condensation ^58^, and synapsin phases that cluster vesicles may be rapidly dispersed by CamKII activation ^7^. Our work uncovers a regulatory pathway that induces structural active zone plasticity through phase separation. We propose that a fraction of Liprin-α3 is soluble and that phosphorylation by PKC nucleates the transition of Liprin-α3 into liquid condensates to recruit additional Liprin-α3, RIM, Munc13-1, and possibly other active zone proteins. This allows addition of secretory machinery to the membrane to enhance release.

Modulation of Liprin-α3 phase separation by PKC complements the other presynaptic mechanisms for PLC/PKC-triggered potentiation, including those mediated by Munc13 ^38^, Munc18 ^39^ and Synaptotagmin-1 ^40^, further supporting the involvement of multiple parallel mechanisms ^27^.

Finally, the question arises whether Liprin-α3 phase separation may control synapse and active zone formation during development. This appears likely because S760-phosphorylated Liprin-α3 is more prominent early postnatally and blocking phase separation of Liprin-α3 throughout development results in basal defects of active zone structure. Liprin-α liquid condensates may interact with a wide range of synaptic proteins to broadly orchestrate assembly ^5, 12, 13, 21–25^. This may include interactions with vesicles, cytoskeletal elements and trafficking machinery, potentially explaining why some synaptic vesicles are lost in Liprin-α deficient synapses ^9–11, 59, 60^. During development, it is likely that phase separation and recruitment of presynaptic material occur independent of phosphorylation and involve additional proteins. For instance, other Liprin-α isoforms or ELKS, which captures synaptic material and phase separates ^32, 61^, may play active roles. In conclusion, an overarching model arises in which phase transition of presynaptic proteins is essential to recruit and assemble presynaptic material into functional molecular machines.

## Acknowledgements

We thank J. Wang, E. Atwater, M. Sanghvi and M. Han for technical support, Drs. R. Held, C. Tan and N. Nyitrai for help and advice, and all members of the Kaeser laboratory for insightful discussions. We thank Dr. S. Schoch for Liprin-α antibodies, and Drs. M. Verhage and J. Broeke for the SynapseEM MATLAB macro. This work was supported by grants from the NIH (R01NS083898 and R01MH113349 to PSK, R35GM130386 to T.K.), the Lefler Foundation (to PSK), the Armenise Harvard Foundation (to PSK), a grant from the Novo Nordisk Foundation/Danish Technical University (NNF16OC0022166 to T.K.), a Biogen Sponsored Research Agreement (to T.K), and fellowships from the Alice and Joseph E. Brooks postdoctoral fund (to JEM), the Croucher foundation (to MYW), Lefler foundation (to MYW) and the NSF (graduate research fellowship DGE1144152 to S.S.H.W.). We acknowledge the Neurobiology Imaging Facility (supported by a P30 Core Center Grant NS072030), and the Electron Microscopy Facility at Harvard Medical School.

## Author Contributions

Conceptualization, J.E-M., M.Y.W. and P.S.K.; Methodology, J.E-M., M.Y.W., G.dN. and T.K.; Investigation, J.E-M., M.Y.W., S.S.W. and G.dN.; Formal Analysis, J.E-M., M.Y.W., S.S.W., G.dN., T.K. and PSK; Writing-Original Draft, J.E-M and P.S.K.; Supervision, P.S.K.; Funding Acquisition P.S.K.

## Materials and methods

### Assessments of droplets in transfected HEK293T cells

HEK293T cells were plated on 0.1 mm thick coverslips and transfected with plasmids expressing proteins of interest under the CMV promoter. 500 ng of DNA per well (1,9 cm^2^) were used for single plasmid transfections. If multiple plasmids were transfected, additional DNA was used at a 1:1 molar ratio. Cultures were fixed in 4% paraformaldehyde 10 – 16 h after transfection. Longer expression times or higher amounts of DNA were avoided to limit protein aggregation. Drugs were added 15 min before cells were fixed at the following concentrations: forskolin (10 µM, Sigma), phorbol 12-myristate 13-acetate (PMA, 1 µM, Sigma), caffeine (1 mM, Sigma), H-89 (5 µM, Abcam), bisindolylmaleimide-I (Bis-I, 0.1 µM, Sigma), KN-93 (1 µM, Abcam) and cells were fixed in the presence of drugs. When non-fluorescently-tagged proteins were expressed, staining with primary (rabbit anti Liprin-α1 (A121), -α2 (A13), -α3 (A115) and - α4 (A2) 1:250; gifts from S. Schoch ^35^) and 488 Alexa-conjugated secondary antibodies (overnight at 4°C in both cases) was performed. Images were acquired with a Leica SP8 Confocal/STED 3X microscope, using an oil-immersion 63X objective. For single protein expression, quantification was done manually, including only spherical condensates or rings of >1 µm in diameter. To quantify the amount and size of protein structures created by Liprin-α3, RIM1α and Munc13-1, the “Analyze particles” plug-in (Fiji) was used with automatic thresholding of the Munc13-1 channel and a minimum diameter of 1 µm. In all experiments comparing different proteins or treatments, the experimenter was blind to the condition throughout data acquisition and analyses. For Fluorescence Recovery After Photobleaching (FRAP), HEK293T cells were plated on 35-mm plastic dishes containing 0.15 mm thick coverslips. 12 – 15 h after transfection and 10 min after PMA addition, the dishes were transferred to the microscope stage and single droplets or peripheral condensates were photobleached using a 405 nm wavelength laser followed by image acquisition at a 1 (Fig. 1) or 3 (Fig. 3) Hz sampling frequency in confocal mode. HEK293T cells were kept in the tissue culture medium containing 1 µM PMA and imaged at room temperature within 1 h of PMA addition. Regions of interest were drawn over pre-bleached structures and the percentage of intensity recovered was plotted as a function of time. t_1/2 recovery_ was calculated as the time it takes for fluorescence to reach 50% of the maximum recovery after bleaching. Images were acquired using a Leica SP8 Confocal/STED 3X microscope, using an oil-immersion 63X objective. The following N-terminally tagged (unless noted otherwise) plasmids were used: pCMV HA-Liprin-α1 (p462), pCMV HA-Liprin-α2 (p463), pCMV HA-Liprin-α3 (p470), pCMV GFP-Liprin-α4 (p466), pCMV Cerulean-Liprin-α3 (p471), pCMV mVenus-Liprin-α3 (p472), pCMV mVenus-Liprin-α3 Y648A+S650A+S651A (p516), pCMV mVenus-Liprin-α3 S751A (p499), pCMV mVenus-Liprin-α3 S760A (p500), pCMV mVenus-Liprin-α3 S760G (p507), pCMV mVenus-Liprin-α3 S760E (p503), pCMV mVenus-Liprin-α3 S763A (p510) and pCMV mVenus-Liprin-α3 S764A (p514), pCMV RIM1α-mVenus (p587; tag placed before the C2B domain, which does not interfere with protein function ^33, 43^), and pcDNA Munc13-1-tdTomato (p888; tag placed at the C-terminus).

### Expression and purification of GST-Liprin-α3 proteins

GST-tagged fusion proteins were generated, expressed and purified according to standard procedures and as described ^32^. Briefly, proteins were expressed at 20°C in E. coli BL21 cells after induction with 0.05 mM isopropyl b-D-1-thiogalactopyranoside for 20 h, and pelleted by centrifugation (45 min on 3,500 x g). For purification of GST-fusion proteins, bacterial pellets were resuspended and lysed for 30 min in PBS buffer supplemented with 0.5 mg/mL lysozyme, 0.5 mM EDTA, and a protease inhibitor cocktail, followed by brief sonication and centrifugation (45 min on 11,200 x g). Next, bacterial supernatants were incubated with glutathione-Sepharose resin (GE Healthcare) for 1.5 h at 4°C with gentle rotation, washed three times in PBS and stored until further use (for no more than 5 d after purification). All steps after protein induction were conducted at 4°C using ice-cold solutions. Protein concentrations were estimated in SDS-gel electrophoresis and Coomassie staining using increasing BSA concentrations as reference. The following GST-tagged proteins were produced from pGEX-KG2 constructs: pGEX Liprin-α3 1 – 188 (p567), pGEX Liprin-α3 189 – 576 (p568), pGEX Liprin-α3 577 - 790, (p566) and pGEX Liprin-α3 791 – 1192 (p570). Amino acid numbering follows NM_001270985.2.

### In-vitro phosphorylation of Liprin-α3 domains

40 μg of fusion proteins bound to glutathione beads were incubated for 30 min in 200 μL of PKC reaction buffer (20 mM HEPES, 10 mM MgCl_2_, 1.67 mM CaCl_2_, 150 mM NaCl_2_, 1 mM DTT) with 0.25 ng/μl PKC (Promega, V526A), 1 μM PMA, 1 μM Phosphatidyl Serine (Sigma, P7769) and 200 μM ATP (Sigma, A2383). For experiments in which phosphorylation was detected by autoradiography, 10 µCi ^32^P-γ-ATP (Perkin Elmer) was added to the PKC reaction mix and incubated for an additional 1 hr at 30 °C, followed by gel electrophoresis. For mass spectrometric analysis, the phosphorylated GST Liprin-α3 577 - 790 protein was isolated by SDS gel electrophoresis, Coomassie blue staining and cutting out of the protein band after the initial PKC reaction. The sample was processed by the HMS Taplin Mass Spectrometry Facility for identification of phosphorylated amino acid residues.

### Generation of custom antibodies

Custom antibodies (A231, A232 and 247) were generated using procedures as described ^34^. Phospho-specific Liprin-α3 antibodies were generated using keyhole lympet hemocyanin (KLH) conjugated CKAPKRK(pSer760)IKSSIGR or CAPKRKSIKS(pSer764)IGRL. GST-Liprin-α3 188- 576 (p568) was expressed in and purified from BL21 bacteria. Peptides and GST-fusion proteins were injected into rabbits whose sera had been pre-screened to prevent non-specific antibody signal, boosters were given every 2 weeks and bleeds were collected every 3 weeks. Sera with strong Liprin specificity in western blotting were processed by affinity purification ^34^.

### Western blotting

Samples were prepared in SDS sample buffer as described ^32^, run on SDS-PAGE gels and transferred to nitrocellulose membranes at 4 °C for 6.5 hr in buffer containing (per L) 200 mL methanol, 14 g glycine and 6 g Tris, followed by a 1 h block at room temperature in saline buffer with 10% non-fat milk powder and 5% normal goat serum. Primary antibodies were incubated overnight at 4 °C in saline buffer with 5% milk and 2.5% goat serum, followed by 1 h incubation at room temperature with horseradish peroxidase-conjugated secondary antibodies prior to visualization of the protein bands. Primary antibodies used: rabbit anti Liprin-α1 (A121, 1:500), Liprin-α2 (A13, 1:500), Liprin-α3 (A115, 1:500) and Liprin-α4 (A2, 1:500) were gifts from S. Schoch ^35^; rabbit anti phospho-760 Liprin-α3 (generated for this study; A231; 1:1000) and phospho-764 Liprin-α3 (generated for this study; 1:1000); mouse anti HA (A12, 1:500; RRID: AB_2565006); mouse anti Synaptophysin (A100, 1: 5000; RRID:AB_887824) and mouse anti Synapsin-1 (A57, 1:5000; RRID: AB_2617071). Three 5 min washes were performed between steps.

### Neuronal cultures and production of lentiviruses

Primary hippocampal cultures were prepared as described ^16, 32–34^. Briefly, newborn (P0-P1) pups were anesthetized on ice slurry prior to hippocampal dissection. Hippocampi were digested and dissociated, and neurons were plated onto glass coverslips in Plating Medium composed of Mimimum Essential Medium (MEM) supplemented with 0.5% glucose, 0.02% NaHCO3, 0.1 mg/mL transferrin, 10% Fetal Select bovine serum, 2mML-glutamine, and 25 mg/mL insulin. 24 h after plating, Plating Medium was exchanged with Growth Medium composed of MEM with 0.5% glucose, 0.02% NaHCO3, 0.1 mg/mL transferrin, 5% Fetal Select bovine serum (Atlas Biologicals FS-0500-AD), 2% B-27 supplement, and 0.5 mM L-glutamine. At DIV2-3, 4 mM Cytosine b-D-arabinofuranoside (AraC) was added. Cultures were kept in a 37 °C tissue culture incubator until DIV15 – 17. Lentiviruses were produced in HEK293T cells maintained in DMEM supplemented with 10% fetal bovine serum and 1% penicillin/streptomycin. HEK293T cells were transfected using the calcium phosphate method with the lentiviral packaging plasmids REV, RRE and VSV-G and a separate plasmid encoding the protein of interest, at a molar ratio 1:1:1:1. 24 h after transfection, the medium was changed to neuronal growth medium and, 18 - 30 later the supernatant was used for immediate transduction. Neuronal cultures were infected 4 - 5 d after plating with lentiviruses expressing GFP-Cre or an inactive variant of GFP-Cre expressed under the human Synapsin promotor ^62^. For rescue, cultures were infected at DIV1 – 2 with a lentivirus expressing Liprin-α3 or Liprin-α3^SG^, or an empty lentivirus as control. pFSW HA-Liprin-α3 S760G was generated for this study; pFSW control (p008) and pFSW HA-Liprin-α3 (p526) were previously described ^16^. For PMA experiments, PMA was added 15 - 20 min before fixation to a final dilution of 1 µM (from a 1 mM stock diluted in DMSO), and neurons were washed and fixed or recorded in the presence of the drug. – PMA controls were incubated in the same amount of DMSO.

### Immunofluorescence staining and confocal microscopy of neurons

Neurons grown on #1.5 (for STED) or # 1.0 (confocal) glass coverslips were fixed in 4% paraformaldehyde for 10 min at DIV15-17, blocked and permeabilized in blocking solution (3% BSA/0.1% Triton X-100/PBS) for 1 h, incubated overnight with primary antibodies followed by overnight incubation with Alexa-conjugated secondaries (Thermo Fisher), and mounted onto glass slides. Antibodies were diluted in blocking solution. For STED imaging, coverslips were additionally post-fixed in 4% paraformaldehyde for 10 min. Three 5 min washes with PBS were performed between steps. All steps were performed at room temperature except for antibody incubations (4 °C). Primary antibodies used: mouse anti Bassoon (A85, 1:50; RRID:AB_11181058), rabbit anti Liprin-α2 (A13, 1:250; gift from S. Schoch ^35^) and -α3 (A115, 1:250; gift from S. Schoch ^35^), rabbit anti RIM (A58, 1:500, RRID: AB_887774), mouse anti PSD-95 (A149, 1:500; RRID: AB_10698024), mouse anti Gephyrin (A8, 1:500; RRID:AB_2232546), rabbit anti Synapsin-1 (A30, 1:500; RRID:AB_2200097), mouse anti Synapsin-1 (A57, 1:500; RRID: AB_2617071), guinea pig anti Synaptophysin (A106, 1:500; RRID: AB_1210382), rabbit anti RIM-BP2 (A126, 1:500; RRID: AB_2619739), rabbit anti Munc13-1 (A72, 1:500; RRID: AB_887733), rabbit anti Ca_V_2.1 (A46, 1:500; RRID: AB_2619841), mouse anti HA (A12, 1:500; RRID: AB_2565006), mouse anti MAP2 (A108, 1:500; RRID: AB_477193), rabbit anti MAP2 (A139, 1:500; RRID: AB_2138183), mouse anti GluA1 (A82; 1:100; RRID:AB_2113443). Confocal images were taken on an Olympus FV1200 confocal microscope equipped with a 60X oil immersion objective or a Leica SP8 Confocal/STED 3X microscope with a 63X oil immersion objective. Images of experiments with multiple groups were acquired within a single session per culture and identical settings for each condition were used within an imaging session. For quantitative analyses of synaptic protein levels, the synaptic vesicle marker signal was used to define puncta as ROIs, and the average intensity within ROIs was quantified after local background was subtracted using the “rolling average” ImageJ plugin (diameter = 1.4µm). Data was plotted normalized to the average intensity of the control group (control^L23^) per culture. For co-localization analyses, the “Coloc 2” imageJ plugin was used following default thresholding. For example images, brightness and contrast were linearly adjusted equally between groups and interpolated to meet publication criteria.

### STED Imaging of synapses

Images were acquired with a Leica SP8 Confocal/STED 3X microscope equipped with an oil-immersion 100X 1.44-N.A objective, white lasers, STED gated detectors, and 592 nm and 660 nm depletion lasers as described ^16, 32–34^. Synapse-rich areas were selected and were scanned at 22.5 nm per pixel. Triple color sequential confocal scans were followed by dual-color sequential STED scans. Identical settings were applied to all samples within an experiment. For quantification, side-view synapses (selected while blind to the protein of interest) were defined as synapses that contained a vesicle cluster (imaged in confocal mode, >300 nm wide) with an elongated Bassoon, Gephyrin or PSD-95 (active zone or postsynaptic density markers, respectively, imaged by STED) structure along the edge of the vesicle cluster ^16, 32–34^. A 1 μm-long, 250-nm-wide profile was selected perpendicular to the active zone/postsynaptic density marker and across its center. The intensity profile was then obtained for markers and for the protein of interest. Peak levels of the protein of interest were measured as the maximum intensity of the line profile within 100 nm of the active zone/postsynaptic density marker peaks (estimated active zone area based on ^16^) after applying a 5-pixel rolled average. Only for representative images, a smooth filter was added and brightness and contrast were linearly adjusted using ImageJ. Equal adjustments were performed for all images within a given experiment. Finally, images were interpolated to match publication standards. Quantitative analyses were performed on original images without any processing, and all data were acquired and analyzed by an experimenter blind to genotype and/or condition.

### Mouse lines

Liprin-α2 (*Ppfia2*) mutant mice were acquired from MRC Harwell (C57BL/6N-Ppfia2<tm1a(EUCOMM)Hmgu>/H) ^63^. The mice were generated by homologous recombination and first were crossed to Flp-expressing mice ^64^ to remove the LacZ/Neomycin cassette to generate the conditional allele, which contains loxP sites flanking exon 14. Conditional Liprin-α2 mice were kept as homozygotes, and genotyped using oligonucleotide primers GCCTCTTAACATTCACTGTACC and CCAGTGTGTACTGGAGACAAGC for the wild-type allele (336 band), and GCCTCTTAACATTCACTGTACC and CTGCGACTATAGAGATATCAACC for the floxed allele (517 band). To generate Liprin-α2/α3 double mutant mice, conditional Liprin-α2 knockout mice were crossed to previously described constitutive Liprin-α3 mice that were generated by CRISPR/Cas9-mediated genome editing ^16^. The line was maintain using intercrosses between Liprin-α2^f/f^/Liprin-α3^-/-^ and Liprin-α2^f/f^/Liprin-α3^+/-^ mice. For experiments, hippocampal neurons cultured from individual P0 Liprin-α2^f/f^/Liprin-α3^-/-^ pups were infected with lentivirus expressing Cre recombinase (to generate KO^L23^ neurons) and compared to Liprin-α2^f/f^ x Liprin-α3^+/-^ littermates infected with lentiviruses that express a truncated, inactive mutant of Cre (to generate control^L23^ neurons) ^62^, both expressed via a human synapsin promoter. For rescue experiments comparing Liprin-α3 with Liprin-α3^SG^, the genotype of the breeders was Liprin-α2^f/f^/Liprin-α3^-/-^ and neurons were cultured from pooled hippocampi from multiple pups of the same litter, followed by addition of rescue virus and of lentivirus expressing Cre recombinase as described under neuronal cultures. All animal experiments were approved by the Harvard University Animal Care and Use Committee.

### Electron microscopy of cultured neurons

Electron microscopy was performed as described ^19, 34^. Briefly, neurons grown on 0.12-mm-thick carbon-coated sapphire coverslips were transferred to extracellular solution containing (in mM) 140 NaCl, 5 KCl, 2 CaCl2, 2 MgCl2, 10 glucose, 10 Hepes (pH 7.4, ∼310 mOsm) and subsequently frozen with a Leica EM ICE high-pressure freezer at DIV15-17. After freeze substitution (in acetone containing 1% osmium tetroxide, 1% glutaraldehyde, and 1% H_2_O), samples were embedded in epoxy resin and sectioned at 50 nm with a Leica EM UC7 ultramicrotome. Samples were imaged with a JEOL 1200EX transmission electron microscope equipped with an AMT 2k CCD camera. Images were analyzed using SynapseEM, a MATLAB macro provided by Dr. Matthijs Verhage. Bouton size was calculated from the perimeter of each synapse. Docked vesicles were defined as vesicles touching the presynaptic plasma membrane (with no space between the electrondense vesicular and target membranes) opposed to the PSD. All data were acquired and analyzed by an experimenter blind to the genotype.

### Correlative light-electron microscopy

HEK293T cells were grown on photo etched gridded coverslip and fixed 12 – 16 h after transfection in 2.5% glutaraldehyde, 2% sucrose, 50 mM KCl, 2.5 mM MgCl_2_, 2.5 mM CaCl_2_ and 50 mM cacodylate (pH 7.4) for 2 h 4 °C. A spinning disk confocal microscope (3i, Denver, Colorado) equipped with an oil-immersion 63X 1.4 N.A. objective and 488/561 nm lasers was used for the acquisition of fluorescent images. After image acquisition, samples were stained for 2 h in staining solution I (SSI; consisting of 1% OsO4, 1.25% potassium hexacyanoferrate in 100 mM PIPES, pH 7.4,), followed by staining solution II (prepared by diluting 100 times SSI in 1% tannic acid) for 30 min, and incubated in 1% uranyl acetate overnight. All staining steps were done on ice, the sample was protected from light, and three 5-minute washes with ice-cold mili-Q water were performed between steps. Samples were dehydrated with increasing ethanol concentrations (30%, 50%, 70%, 90%, 100%), followed by two washes in 100% acetone, embedded in epoxy resin, baked at 60 °C for at least 36 h and sectioned at 50 nm with a Leica EM UC7 ultramicrotome. A JEOL 1200EX transmission electron microscope equipped with an AMT 2k CCD camera was used for electron image acquisition. Fluorescent and electron microscopy images were aligned using the BigWarp plugin (ImageJ) using the electron micrograph as a fixed image and different arbitrary references for alignment. As references, cell features such as the nucleus and the plasma membrane and the fluorescent signals were used. Multiple independent alignments using different references were conducted to confirm correct alignment.

### Electrophysiology

Electrophysiological recordings were performed as described before ^19, 34^. DIV15 – 16 neurons were recorded in whole-cell patch-clamp configuration at room temperature in extracellular solution containing (in mM) 140 NaCl, 5 KCl, 1.5 CaCl_2_, 2 MgCl_2_, 10 HEPES (pH 7.4) and 10 Glucose. Glass pipettes were pulled at 2 – 4 MΩ and filled with intracellular solutions containing (in mM) 120 Cs-methanesulfonate, 10 EGTA, 2 MgCl_2_, 10 HEPES-CsOH (pH 7.4), 4 Na_2_-ATP, and 1 Na-GTP for excitatory transmission; and 40 CsCl, 90 K-Gluconate, 1.8 NaCl, 1.7 MgCl_2_, 3.5 KCl, 0.05 EGTA, 10 HEPES, 2 MgATP, 0.4 Na_2_-GTP, 10 phosphocreatine, CsOH (pH 7.4) for inhibitory transmission. For evoked responses, 4 mM QX314-Cl was added to the intracellular solution to block sodium channels. Neurons were clamped at -70 mV for IPSC and AMPAR-EPSC recordings, or +40 mV for NMDA-EPSCs. Series resistance was compensated down to 5 – 5.5 MΩ. Recordings in which the series resistance increased to >15 MΩ before compensation were discarded. mEPSCs, mIPSCs and sucrose-evoked release were measured in extracellular solution supplemented with 1 mM TTX, 50 mM D-AP5 and either 20 mM picrotoxin (for EPSCs) or 20 mM CNQX (for IPSCs). 500 mM hypertonic sucrose was applied for 10 s, and the integral of the first 10 s of the response was used to estimate the RRP. Action potential-evoked responses were elicited by focal bipolar electrical stimulation with an electrode made from Nichrome wire and recorded in extracellular solution supplemented with 20 mM CNQX and either 50 mM D-AP5 (for IPSCs) or 20 mM PTX (for NMDAR-EPSCs). A Multiclamp 700B amplifier and a Digidata 1550 digitizer were used for data acquisition, sampling at 10 kHz and filtering at 2 kHz. Data were analyzed using pClamp. In all experiments, the experimenter was blind to the condition throughout data acquisition and analyses.

### Statistics

Normality and homogeneity of variances were assessed using Shapiro or Levene’s tests, respectively. When test assumptions were met, parametric tests (t-test or one-way ANOVA) were used. Otherwise, the non-parametric tests (Mann-Whitney U or Kruskal-Wallis) were used. For paired pulse ratios, a two-way ANOVA was used. Tukey-Kramer or Holm corrections for multiple testing were applied for parametric and non-parametric post-hoc testing. All data were analyzed by an experimenter blind to the drug condition or genotype. For each dataset, the specific tests used are stated in the figure legend.

## Supplement

**Extended Data Figure 1.**
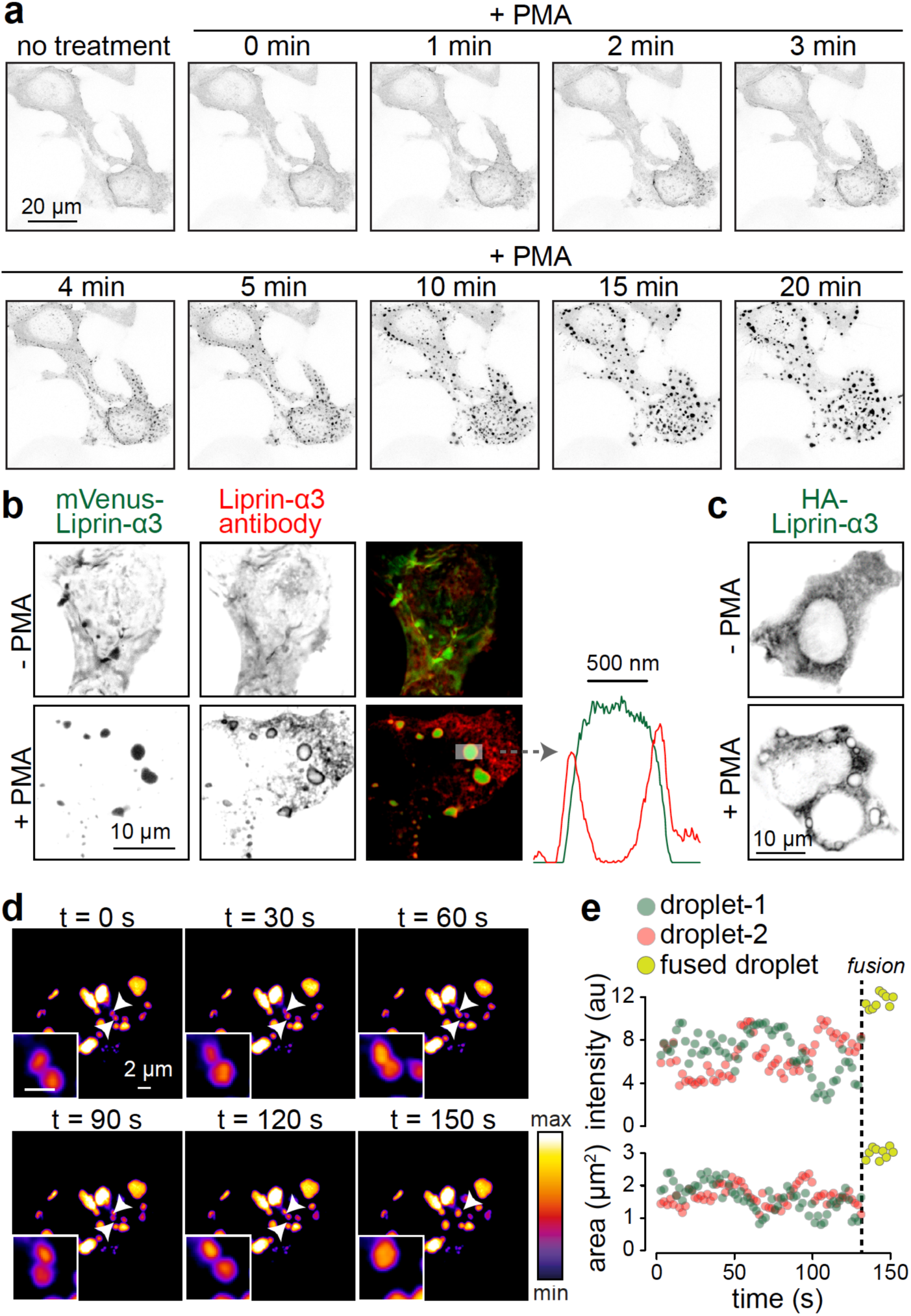
Liprin-α3 forms liquid-liquid phase condensates within minutes after activation of PKC. **(a)** Example confocal time-lapse images of HEK293T cells transfected with mVenus-Liprin-α3 during PMA addition; also see Extended Data Movie 1. **(b)** Example confocal images and line profiles of HEK293T cells transfected with mVenus-Liprin-α3 and immunostained for Liprin-α3. A line profile of a Liprin-α3 condensate is shown on the right. Note that antibody staining produces ring-like shapes around mVenus-Liprin-α3 fluorescence, likely because antibodies do not enter the phase condensates. **(c)** Example of HEK293T cells transfected with HA-Liprin-α3 and immunostained for HA. Note that ring-like structures were only present when PMA was added. **(d, e)** Time-lapse confocal images of two mVenus-Liprin-α3 droplets undergoing a fusion reaction (d), and measurement of the area and intensity of these condensates before and after fusion (e).

**Extended Data Figure 2.**
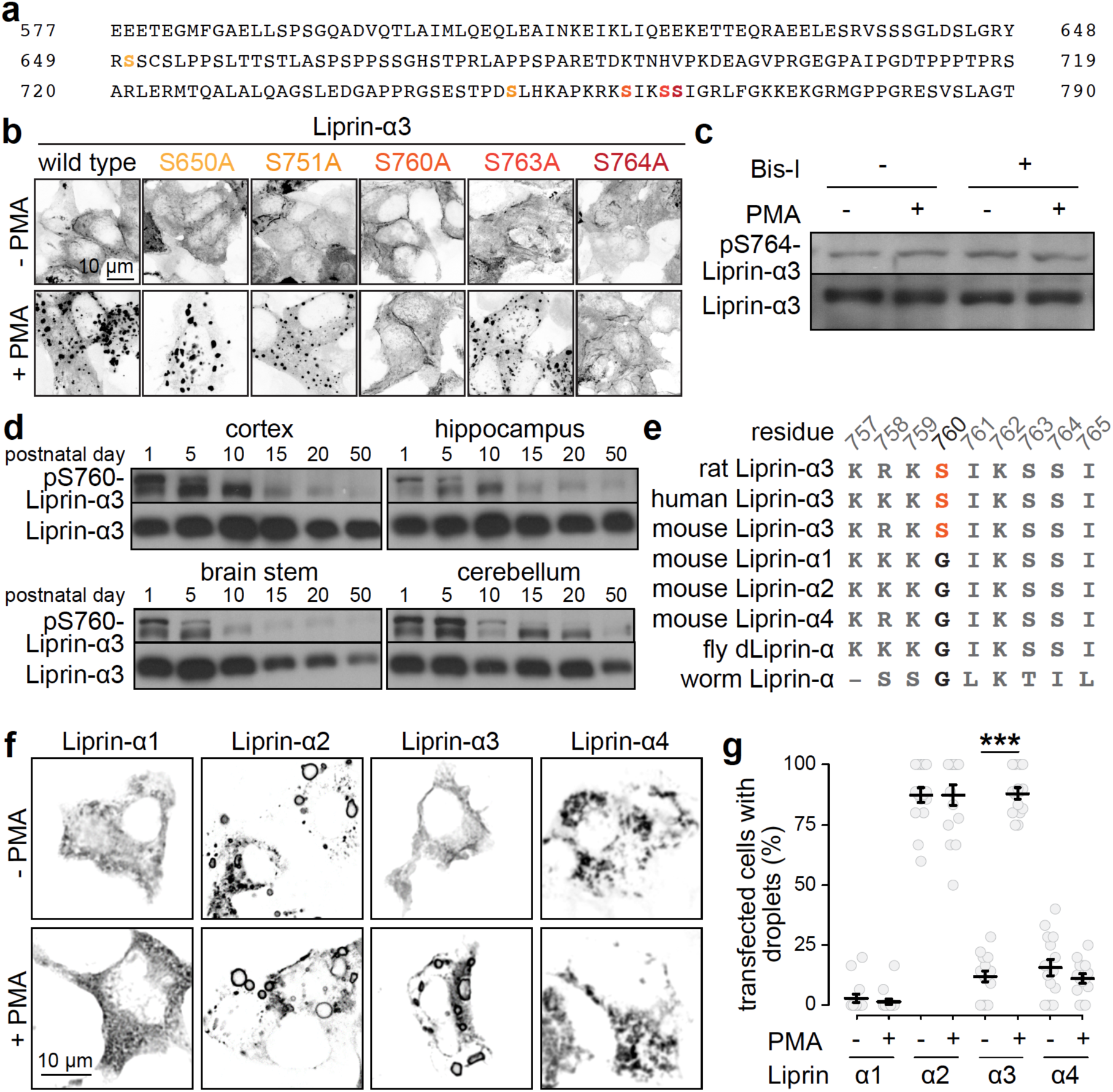
Characterization of Liprin-α3 PKC phosphorylation in vitro and in vivo and assessment of other Liprin-α isoforms. **(a)** Amino acid sequence of the Liprin fragment that was highly phosphorylated. The serines identified by phospho-proteomic analyses are highlighted in a color code repeated in b. **(b)** Confocal images of HEK293T cells transfected with mVenus-tagged Liprin-α3 or mVenus-Liprin-α3 containing point-mutations of each candidate amino acid residues potentially phosphorylated by PKC. Note that S760A and S764A abolish condensate formation upon PMA addition. All constructs contained single point mutations, except for the S650A construct, which also contained Y648A and S651A point mutations. **(c)** Western blot of HEK293T cell lysates transfected with Liprin-α3 showing that activating or blocking PKC phosphorylation does not change the signal detected by phospo-serine-764 specific Liprin-α3 antibodies, and hence serine 764 is unlikely a substrate of PKC. **(d)** Western blots showing the expression profile of phospho-serine-760 Liprin-α3 across brain areas and development. Note the high expression levels during early postnatal days and synaptogenesis. **(e)** Amino acid sequences of various Liprin-α isoforms around serine-760. The PKC phosphorylation site is conserved in Liprin-α3 among vertebrates, but absent in Liprin-α1, -α2 and -α4 and in the single Liprin-α proteins expressed in *C. elegans* and *D. melanogaster*. **(f, g)** Example confocal images (f) and quantification (g) of HEK293T cells transfected with Liprin-α1, -α2, -α3 or -α4 and immunostained for the respective Liprin isoform with or without PMA. Of note, only the PKC-phosphorylatable Liprin-α3 forms ring-like structures, indicative of phase condensates, as a function of the presence of PMA, while Liprin-α2 forms them constitutively. Liprin-α1 and Liprin-α4 do not frequently form such structures. N = 15 images/3 independent transfections per condition. Data in g are mean ± SEM, *** p < 0.001 as assessed by Kruskal-Wallis test with post-hoc Holm tests against the corresponding - PMA control.

**Extended Data Figure 3.**
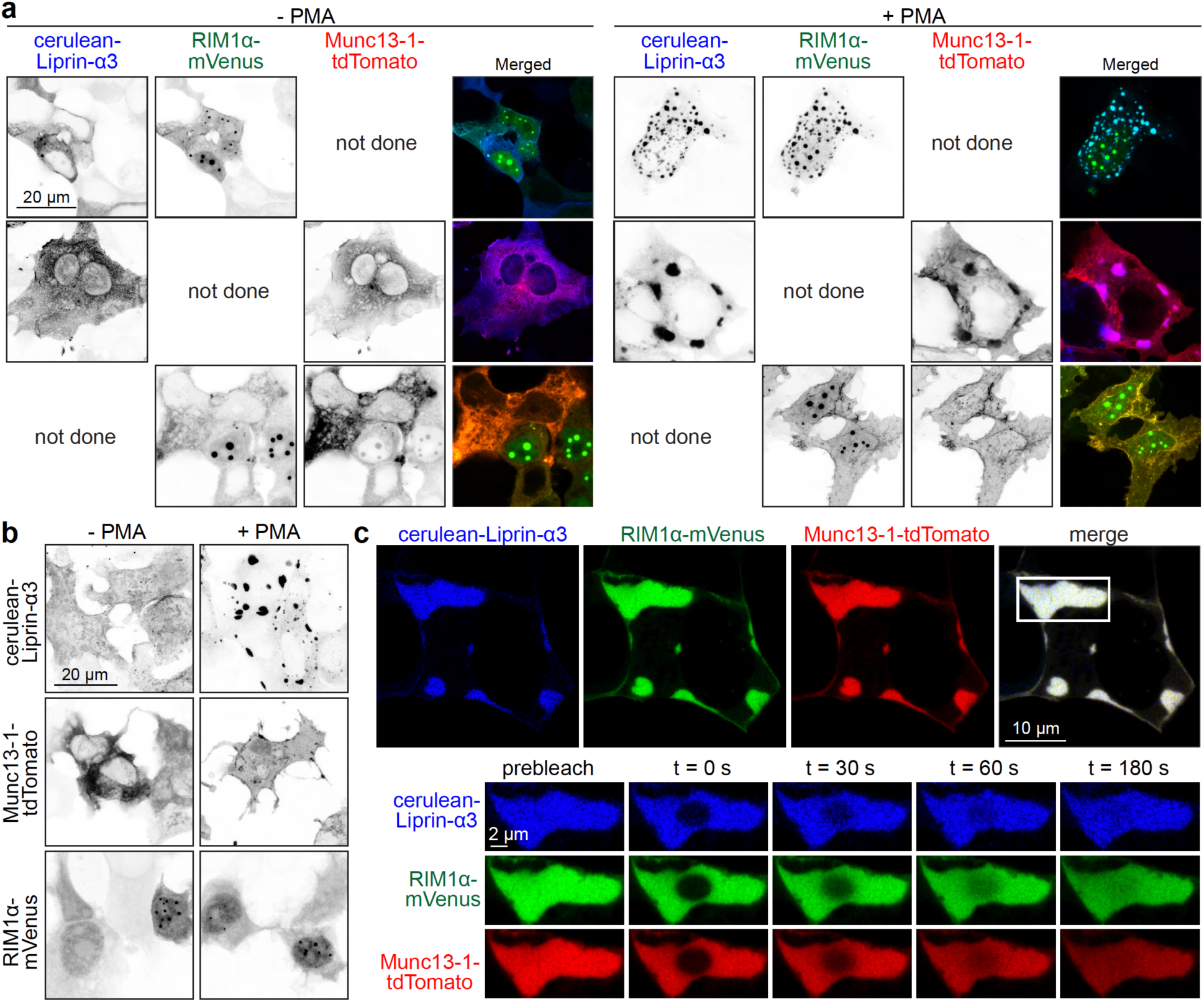
Properties of droplets formed between Liprin-α3, RIM1 and Munc13-1. **(a, b)** Example confocal images of HEK293T cells transfected with combinations of two cDNAs of cerulean-Liprin-α3, RIM1α-mVenus and Munc13-1-tdTomato (a) or with only one cDNA (b) in the presence or absence of PMA. Note that PMA only increases formation of large droplet-like condensates when Liprin-α3 is expressed. **(c)** Example FRAP experiment of a membrane-proximal condensate containing cerulean-Liprin-α3, RIM1α-mVenus and Munc13-1-tdTomato in transfected HEK293T cells in which only the center of the large condensate was photo-bleached. Note fast recovery of all three proteins, indicative of active internal protein rearrangement.

**Extended Data Figure 4.**
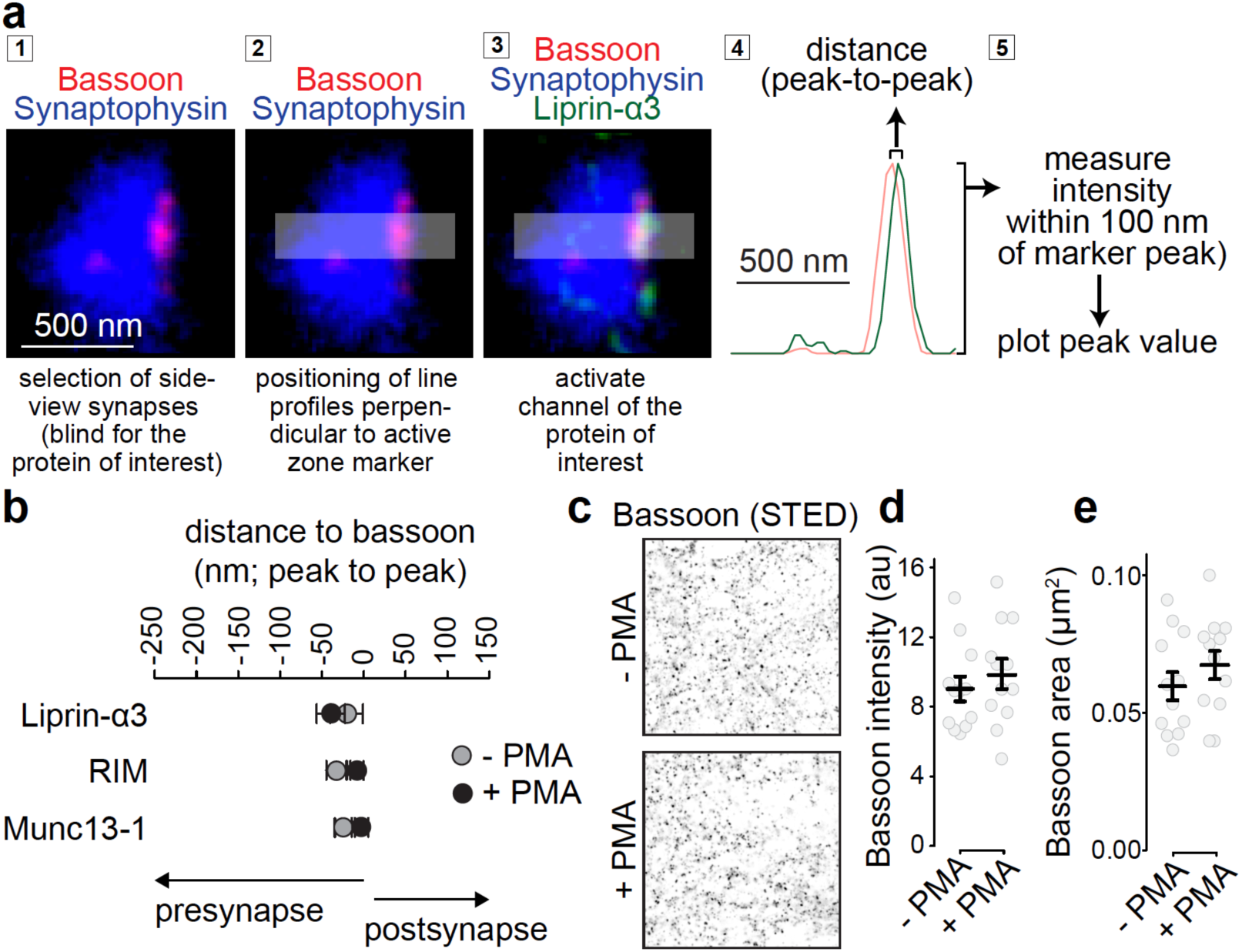
Workflow for STED side-view synapse analysis and peak position of active zone proteins. **(a)** Data analyses workflow for STED side-view synapses, showing an example STED image of a wild type side-view synapse immunostained for Bassoon and Liprin-α3 (imaged in STED mode) and Synaptophysin (imaged in confocal mode). **(b)** Quantification of the average distance of the peak of Liprin-α3, RIM and Munc13-1 of the experiment shown in Figs. 3i-3j to the peak of Bassoon. Liprin-α3: N = 71 synapses/3 independent cultures (- PMA) and 63/3 (+ PMA); RIM: N = 55/3 (-PMA) and 54/3 (+ PMA); Munc13-1 N = 46/3 (- PMA) and 44/3 (+ PMA). **(c-e)** Example Bassoon images (c) and quantification of the average intensity (d) and size (e) of Bassoon objects detected using automatic two-dimensional segmentation (size filter of 0.04-0.4 μm^2^ without considering the shape or orientation of the signal), N = 12 images/3 cultures per condition. Data are shown as mean ± SEM, and Mann-Whitney rank sum test (b) or t-tests (d, e) were used to assess significance.

**Extended Data Figure 5.**
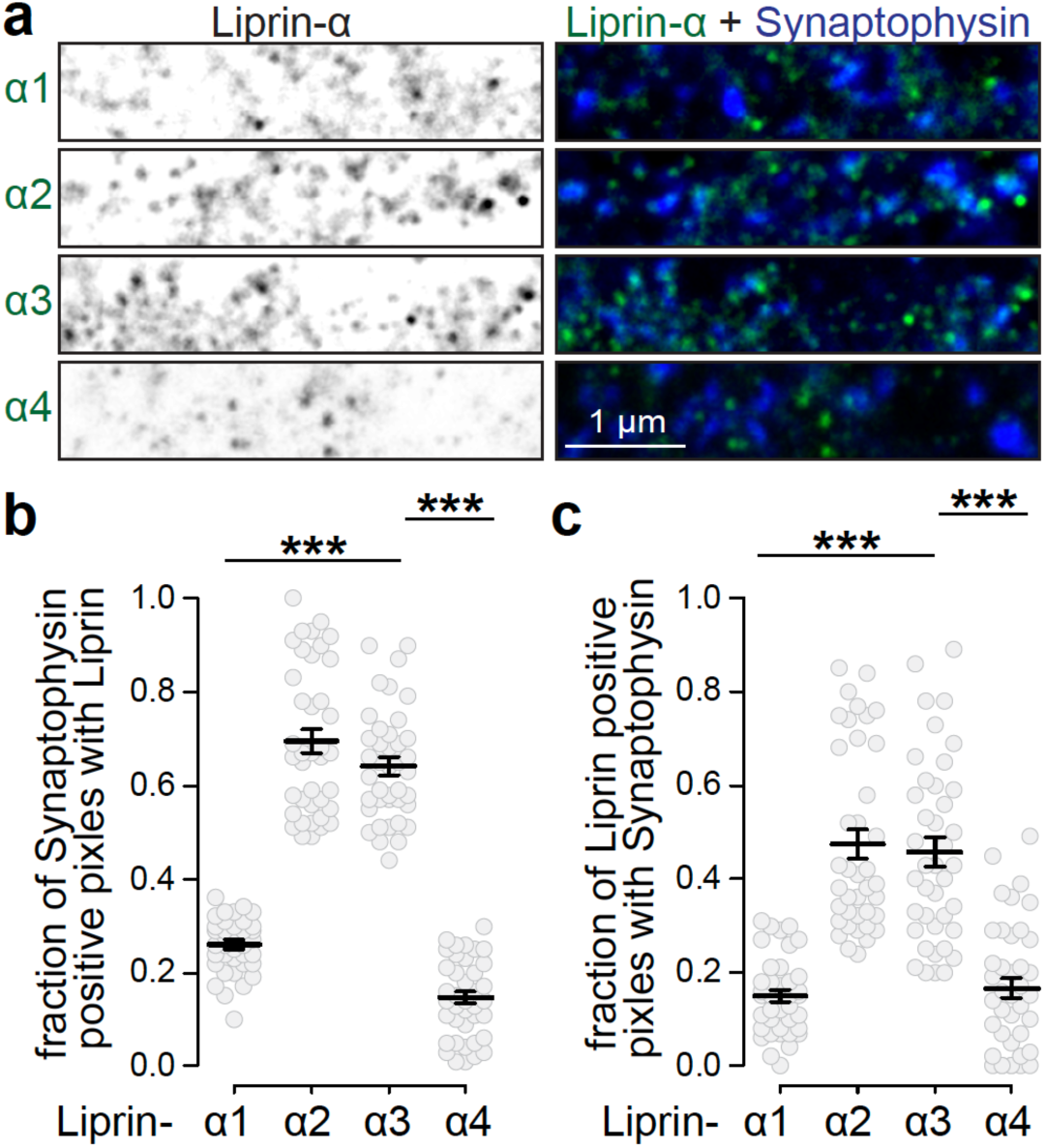
Synaptic expression of Liprin-α proteins. **(a)** Confocal images of wild type mouse hippocampal cultured neurons stained for Synaptophysin and Liprin-α1, -α2, -α3 or -α4. **(b, c)** Mander’s correlation for the fraction of Synaptophysin pixels positive for Liprin-α1, -α2, -α3 or -α4 (b), and vice versa (c). N = 39 images/3 independent cultures for Liprin-α1, -α2; N = 40/3 for Liprin-α3, -α4. Data are shown as mean ± SEM, *** p < 0.001 assessed by Kruskal-Wallis test followed by posthoc Holm comparison against Liprin-α3.

**Extended Data Figure 6.**
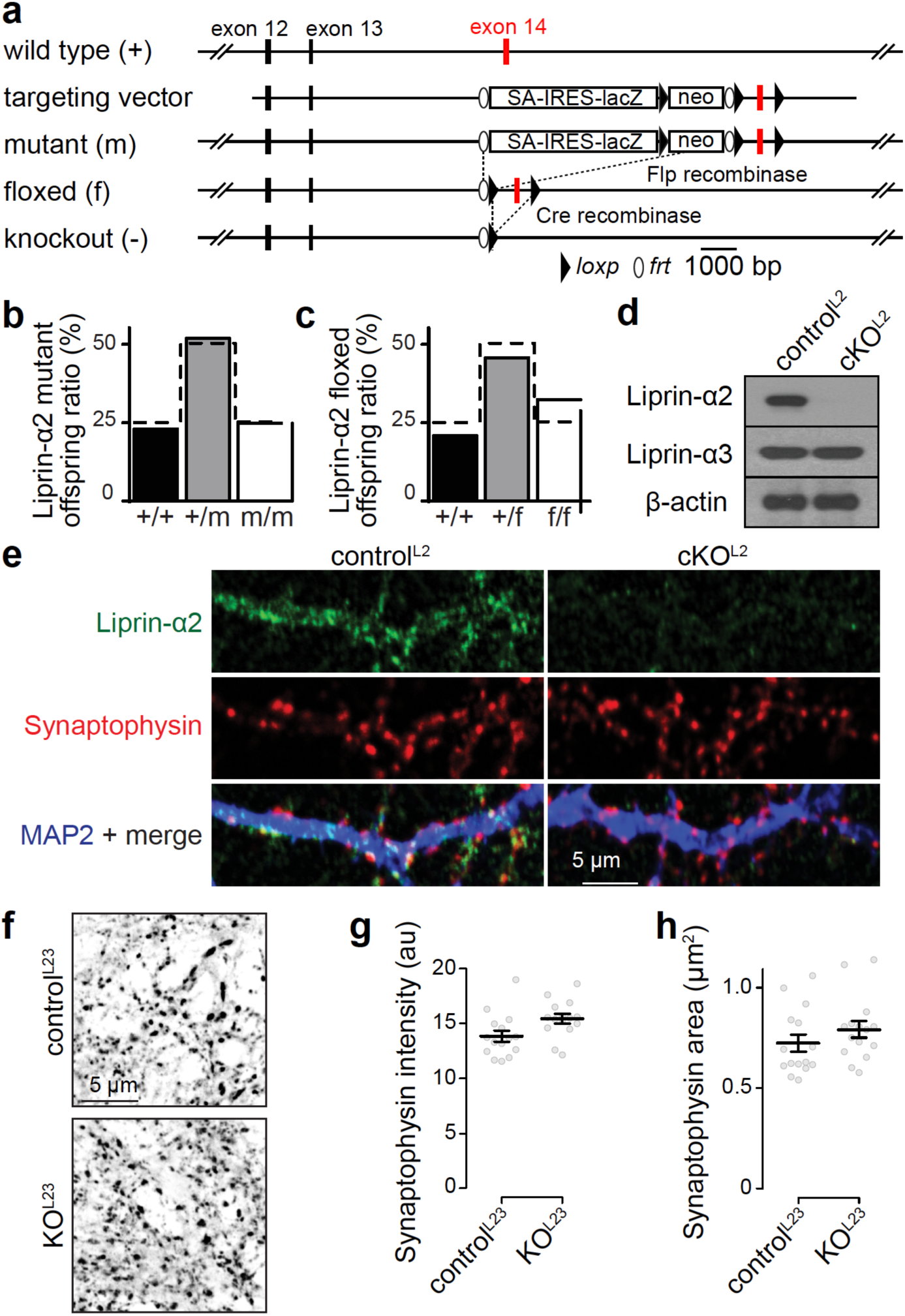
Generation of conditional Liprin-α2 knockout mice. **(a)** Diagram outlining the gene targeting experiment ^63^. The targeting vector contained exon 14 flanked by *loxP* sites, and splice-acceptor/lacZ and neomycin resistance cassettes flanked by *frt* sites. Homologous recombination in embryonic stem cells resulted in the Liprin-α2 mutant allele, and breeding to flp-transgenic mice ^64^ was used to produce the floxed allele. Cre recombinase can then be used to generate the knockout allele. **(b, c)** Survival ratios of the original mutant allele (b) and floxed (c) alleles in a total of ten litters of mice per line. **(d, e)** Western blot (d) and immunostaining (d) of cultured neurons from Liprin-α2 floxed mice that were infected with lentiviruses that express Cre recombinase (to generate cKO^L2^ neurons) or with lentiviruses that express a recombination deficient truncation of Cre (to generate control^L2^ neurons). Liprin-α2 was efficiently removed upon Cre recombination. **(f-h)** Example Synaptophysin images (f) and quantification of the average intensity (g) and size (h) of Synaptophysin objects detected using automatic two-dimensional segmentation (size filter of 0.5-5 μm^2^ without considering the shape or orientation of the signal), N = 15 images/3 cultures per condition. Data are shown as mean ± SEM, no significant differences were observed as tested by a t-test (g) or a Mann-Whitney rank sum test (h).

**Extended Data Figure 7.**
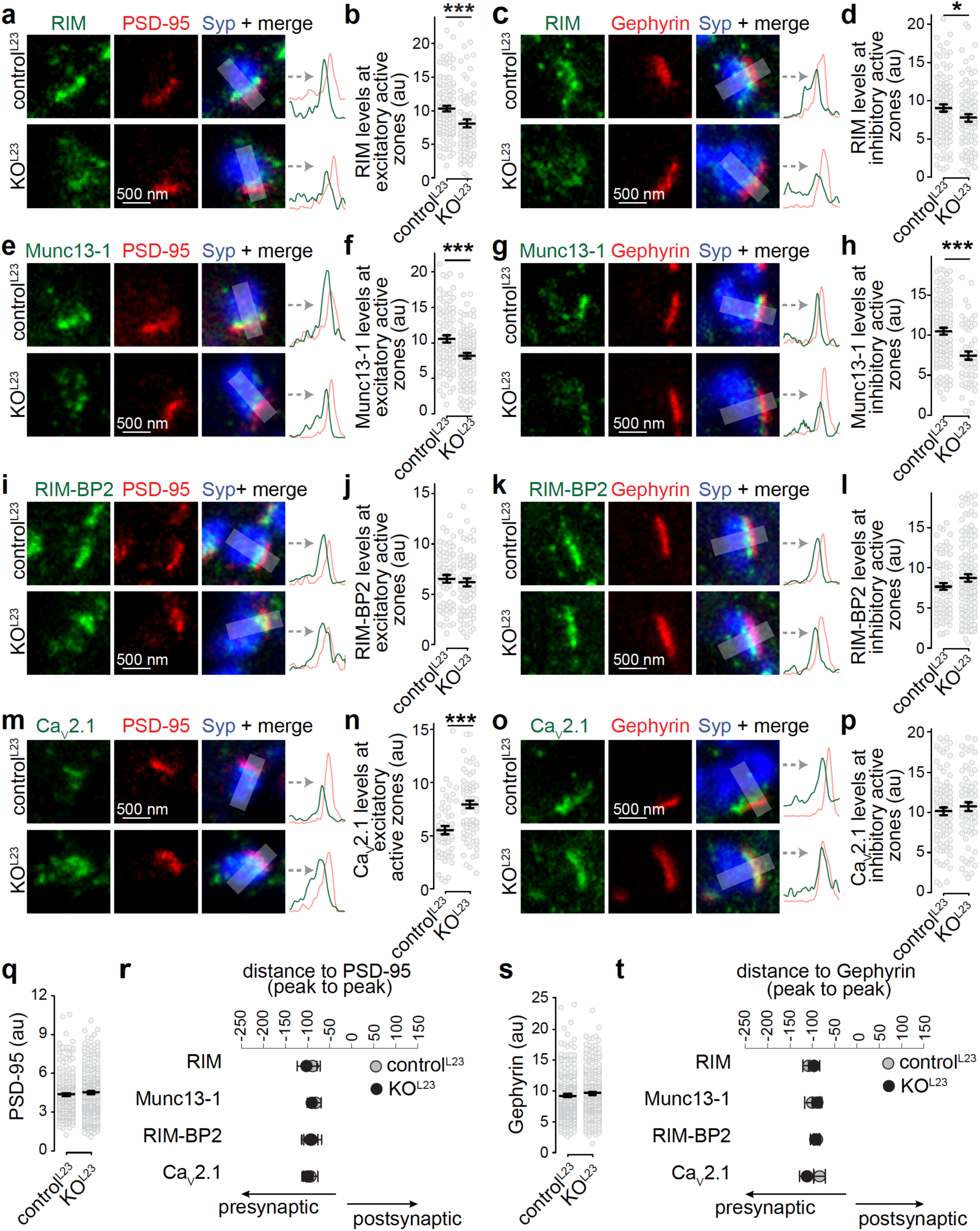
Altered active zone composition after ablation of Liprin-α2 and Liprin-α3. **(a-d)** Example STED images and line profiles (a, c) and quantification of peak intensities (b, d) of RIM at Synaptophysin (Syp) positive excitatory side-view synapses identified via PSD-95 labeling (a, b) or inhibitory side-view synapses identified via Gephyrin labeling (c, d), b: control^L23^: N = 96 synapses/3 independent cultures, KO^L23^: N = 69/3, d: N = 91/3, KO^L23^: N = 87/3. **(e-p)** Same as a-d, but for Munc13-1 (e-h), RIM-BP2 (i-l) and Ca_V_2.1 (m-p). Munc13-1, f: control^L23^: N = 79/3, KO^L23^: N = 99/3, h: control^L23^: N = 102/3, KO^L23^: N = 54/3; RIM-BP2, j: control^L23^: N = 58/3, KO^L23^: N = 68/3, l: control^L23^: N = 73/3, KO^L23^: N = 116/3; Ca_V_2.1, n: control^L23^: N = 53/3, KO^L23^: N = 73/3, p: control^L23^: N = 87/3, KO^L23^: N = 66/3. **(q)** Quantification of the peak intensity of PSD-95 in all line scans analyzed, control^L23^ N = 286/3, KO^L23^ = 309/3. **(r)** Quantification of the average distance of peaks of RIM, Munc13, RIM-BP2 and Ca_V_2.1 to the peak of PSD-95, N as in b, f, j and n. **(s)** Quantification of the peak intensity of Gephyrin in all line scans analyzed, control^L23^ N = 353/3, KO^L23^ = 323/3. **(t)** Quantification of the average distance of peaks of RIM, Munc13, RIM-BP2 and Ca_V_2.1 to the peak of Gephyrin, N as in d, h, l and p. Data are shown as mean ± SEM, * p < 0.05, *** p < 0.001 analyzed by Mann-Whitney rank-sum tests (b, d, h, p, q-t) or t-tests (f, j, l, n).

**Extended Data Figure 8.**
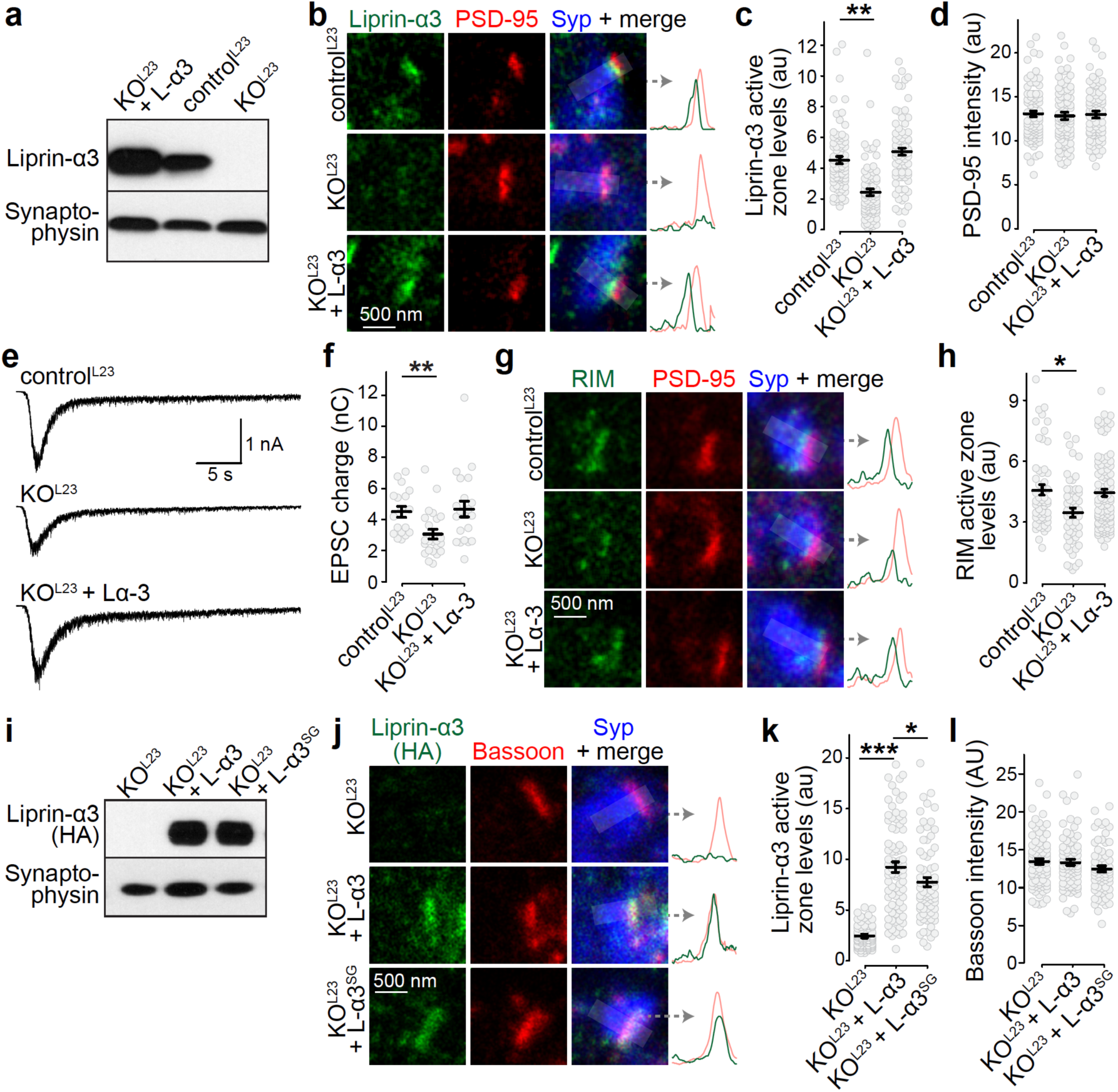
Rescue with Liprin-α3 reverses Liprin-α2/α3 knockout phenotypes. **(a)** Western blot of whole cell lysates of control ^L23^, KO^L23^ and KO^L23^ rescued with Liprin-α3 neuronal cultures. **(b-d)** Example STED images with intensity profiles (b) and quantification (c, d) of the peak intensities of Liprin-α3 and PSD-95 in side-view synapses. control^L23^: N = 83 synapses/3 independent cultures, KO^L23^: N = 82/3, KO^L23^ + L-α3: N = 77/3. **(e, f)** Example traces (e) and quantification (f) of EPSC charge in response to a local 10 s puff of 500 mOsm sucrose to estimate the RRP, control^L23^: N = 18 cells/3 independent cultures, KO^L23^: N = 24/3, KO^L23^ + L-α3: N = 21/3. **(g, h)** Example STED images with intensity profiles (g) and quantification (h) of the peak intensity of RIM at the active zone of side-view synapses, control^L23^: N = 54/3, KO^L23^: N = 54/3, KO^L23^ + L-α3: N = 97/3. **(i)** Western blot of whole cell lysates of KO^L23^ and KO^L23^ rescued with Liprin-α3 or Liprin-α3^SG^ neuronal cultures. An antibody against the HA tag was used for detection. **(j-l)** Representative STED images with intensity profiles (j) and quantification (k, l) of the peak intensities of Liprin-α3 and Bassoon in side-view synapses. control^L23^: KO^L23^: N = 81/3, KO^L23^ + L-α3: N = 86/3, KO^L23^ + L-α3^SG^: N = 74/3. All data are shown as mean ± SEM, * p < 0.05, ** p < 0.01, *** p < 0.001 as analyzed by Kruskal-Wallis tests (c, f, h, k) or one-way ANOVA tests (d, l) with posthoc (Holm and Tukey-Kramer, respectively) testing against control^L23^ (c, f, h) or KO^L23^ and KO^L23^ + L-α3 (k).

**Extended Data Figure 9.**
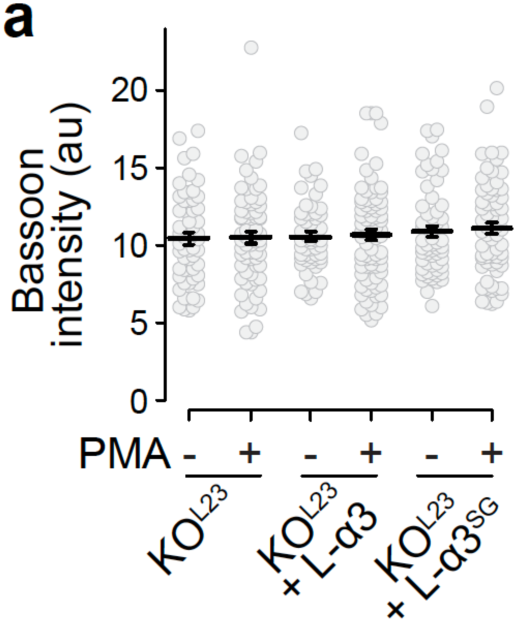
Bassoon levels after re-expression of Liprin-α3 and addition of PMA. **(a)** Quantification of the peak intensity of bassoon (data set from figure 7B). KO^L23^: N = 70 synapses/3 independent cultures (- PMA) and 71/3 (+ PMA), KO^L23^ + L-α3: N = 54/3 (- PMA) and 83/3 (+ PMA); KO^L23^ + L-α3^SG^: N = 63/3 (- PMA) and 73/3 (+ PMA). Data are shown as mean ± SEM and analyzed using a Kruskal-Wallis test.

**Extended Data Movie 1. Liprin-α3 forms droplets within minutes upon addition of PMA** Time-lapse confocal movie showing formation of liquid condensates in HEK293T cells transfected with mVenus-Liprin-α3 upon PMA addition. Black boxes highlight two examples of fusion reactions between condensates. Note that condensates are mobile, in agreement with liquid dynamics, total time of experiment is 20 min compressed to 10 s.

